# Dynamic control of metabolic zonation and liver repair by endothelial cell Wnt2 and Wnt9b revealed by single cell spatial transcriptomics using Molecular Cartography

**DOI:** 10.1101/2022.03.18.484868

**Authors:** Shikai Hu, Silvia Liu, Yu Bian, Minakshi Poddar, Sucha Singh, Catherine Cao, Jackson McGaughey, Aaron Bell, Levi L Blazer, Jarret J Adams, Sachdev S Sidhu, Stephane Angers, Satdarshan P. Monga

**Affiliations:** School of Medicine, Tsinghua University, Beijing, China; Division of Experimental Pathology, Department of Pathology, University of Pittsburgh School of Medicine, Pittsburgh, PA USA; Donnelly Centre, University of Toronto, Toronto, ON, Canada; Leslie Dan Faculty of Pharmacy, University of Toronto, Toronto, ON, Canada; Pittsburgh Liver Research Center, University of Pittsburgh Medical Center and University of Pittsburgh School of Medicine, Pittsburgh, PA USA; Division of Gastroenterology, Hepatology and Nutrition, Department of Medicine, University of Pittsburgh School of Medicine, Pittsburgh, PA USA

## Abstract

The conclusive identity of Wnt proteins regulating liver zonation (LZ) and regeneration (LR) remains unclear despite an undisputed role of β-catenin. Using single-cell analysis of liver cells from various species, a conserved Wnt2 and Wnt9b expression in endothelial cells (ECs) in zone 3 shown to be the major Wnt cell source, was identified. Conditional EC-elimination of Wnt2 and Wnt9b led to perturbation of LZ with not only loss of β-catenin targets in zone 3, but also re-appearance of zone 1 genes in zone 3, unraveling dynamicity as revealed by single-cell spatial transcriptomics using Molecular Cartography. Defective LR observed in the knockouts phenocopied other models of defective hepatic Wnt signaling. Administration of a tetravalent antibody to activate Wnt signaling rescued LZ and LR in the knockouts. Molecular Cartography on the livers of the agonist-treated animal revealed changes in LZ. Administration of the agonist also promoted LR in acetaminophen overdose acute liver failure (ALF) fulfilling an unmet clinical need. Overall, we report an unequivocal role of EC-Wnt2 and Wnt9b in LZ and LR and show the role of Wnt activators as regenerative therapy for ALF.

## Introduction

The Wnt/β-catenin signaling pathway plays fundamental roles in tissue development, homeostasis, repair, regeneration, and tumorigenesis (Clevers et al., 2014; Russell and Monga, 2018). β-catenin transcriptional activity is controlled by Wnt proteins. Once secreted with the help of the cargo protein Wntless (Wls), Wnt proteins bind to their cell surface receptor complex composed of Frizzled (FZD) and low-density lipoprotein receptor-related protein 5/6 (LRP5/6) co-receptors. Receptor activation leads to inactivation of the destruction complex composed of APC-GSK3β-Axin and allows for β-catenin nuclear translocation and target gene expression (Russell and Monga, 2018).

The Wnt/β-catenin signaling pathway and the RSPO-LGR4/5-ZNRF3/RNF43 axis are well known regulators of pericentral gene expression and in turn of metabolic liver zonation (LZ) and also contribute to liver regeneration (LR) after partial hepatectomy (PH) (Benhamouche et al., 2006; Burke et al., 2009; Hu and Monga, 2021; Michalopoulos and Bhushan, 2021; Planas-Paz et al., 2016; Sekine et al., 2006). Mice with liver-specific knockout of β-catenin (β-catenin-LKO) or liver-specific double-knockout of LRP5 and LRP6 (LRP5-6-LDKO) have disrupted pericentral LZ as seen by loss of pericentral expression of glutamine synthetase (GS), CYP2E1 and CYP1A2; and delayed LR due to impaired induction of Cyclin D1 expression (Sekine et al., 2007; Tan et al., 2006; Yang et al., 2014). Among many cell types, hepatic endothelial cells (ECs) are a main source of Wnt ligands in the adult murine liver (Leibing et al., 2018; Preziosi et al., 2018). Mice with EC-specific deletion of Wls (EC-Wls-KO) lack β-catenin-dependent pericentral LZ and have delayed LR, phenocopying β-catenin-LKO and LRP5-6-LDKO (Preziosi et al., 2018).

In mice and humans, there are nineteen Wnt ligands and while several are expressed in various cell types of the liver, their distinct pathophysiological roles *in vivo* are less well understood (Zeng et al., 2007). In murine livers, *Wnt2* and *Wnt9b* are present at the central venous ECs and hepatic sinusoidal ECs near the central vein (Wang et al., 2015). During LR, we previously reported a 45-fold up-regulation in the expression of *Wnt2* and 18-fold up-regulation in the expression of *Wnt9b* in ECs at 12 hours post-PH (Preziosi et al., 2018). Up-regulation of *Wnt2* and *Wnt9b* is also observed in other hepatic regenerating settings (Ding et al., 2010; Zhao et al., 2019). While these correlative studies suggest roles of Wnt2 and Wnt9b in the liver, conclusive proof supporting roles of Wnt2 and Wnt9b in LZ and LR requires genetic evidence.

Mechanisms of LR have been studied for several decades. However, there are no targeted therapies yet available to stimulate LR in the advanced stages of chronic and acute liver diseases. Since the Wnt/β-catenin pathway contributes to both LZ and LR, it is a promising candidate pathway for drug development for regenerative medicine (Alvarado et al., 2016; Fanti et al., 2014). Recently, a water-soluble small molecule F^P+P^-L6^1+3^ (abbreviated FL6.13) was developed through rational design (Tao et al., 2019). FL6.13 is a tailored tetravalent antibody consisting of two pan-FZD paratopes (recognizing FZD1, FZD2, FZD4, FZD5, FZD7 and FZD8) at the N-termini of the Fc and two anti-LRP6 paratopes at their C-termini-one paratope each for the WNT1 (propeller E1-E2) and another for the WNT3A binding site (propeller E3-E4) on LRP6. FL6.13 is thought to promote Frizzled and LRP6 clustering and to stabilize receptor conformations compatible with downstream activation and robust β-catenin transcriptional activity (Tao et al., 2019). FL6.13 also showed strong activity at nanomolar concentrations on both murine and human cell lines, organoids, and *in vivo* (Tao et al., 2019), and thus could be a highly innovative therapeutic modality in cell therapy and tissue repair.

Here, we analyzed cell type-specific expression of Wnt gene transcripts across different species using single-cell RNA (scRNA) sequencing data that allowed us to identify Wnt2 and Wnt9b secreted from ECs. To directly address the requirement of Wnt2 and Wnt9b produced by ECs, we generated Wnt2 and Wnt9b double floxed mice to breed to lymphatic vessel endothelial hyaluronan receptor 1 (Lyve1)-Cre mice and generated EC-Wnt2-KO, EC-Wnt9b-KO, and EC-Wnt2-9b-DKO mice. The EC-Wnt2-9b-DKO mice exhibited perturbed metabolic LZ as revealed using 100-gene single-cell spatial transcriptomics by Molecular Cartography, which was rescued by FL6.13. The EC-Wnt2-9b-DKO mice also showed delayed LR after 70% PH, which was rescued by FL6.13 that also restored Cyclin D1 gene expression. In fact, FL6.13 engages intact FZD and LRP6 receptors on hepatocytes to induce pericentral gene expression ectopically along with cell proliferation program as shown by the 100-gene single-cell spatial analysis. As a result, FL6.13 promoted LR in murine model of surgical resection and rescued delayed acetaminophen (APAP) overdose-induced hyperacute liver injury, which fulfills a major unmet clinical need.

## Results

### Endothelial expression of *Wnt2* and *Wnt9b* is evolutionarily conserved

The Wnt/β-catenin signaling pathway is an evolutionarily conserved pathway with vital cellular functions. Since the Wnt/β-catenin signaling pathway is important for liver function, we sought to query the expression of various Wnt genes in various liver cells through examination of public scRNA sequencing data from different species (Guilliams et al., 2022). In the human dataset, 18 out of 19 WNTs were detected in various hepatic cell types (Fig. S1). ECs were the predominant source of *WNT2, WNT2B*, and *WNT9B*, although signal of *WNT9B* was weak (Fig. 1A, Fig. S1). In monkey, *WNT2, WNT2B, WNT3, WNT9A,* and *WNT9B* were highly expressed among 16 detected WNTs (Fig. 1A, Fig. S2). ECs expressed high levels of *WNT2* and *WNT9B*, while *WNT2B* was mainly expressed by Kupffer cells. In pig, a possible source for xenotransplantation, 9 WNTs were detected (Fig. 1A, Fig. S3). ECs in porcine liver also expressed very high levels of *WNT2* and *WNT9B. WNT2B* was expressed at a very low level. In mice, 15 Wnts were detected of which *Wnt2* and *Wnt9b* were highest in ECs (Fig. 1A, Fig. S4). Even in mice with NAFLD, *Wnt2* and *Wnt9b* were still predominantly expressed in the ECs, suggesting preservation of their fundamental function even in some pathological states (Fig. S5). Altogether, as could be seen from analysis in multiple species (Fig. 1A), ECs are uniformly the predominant source of both *Wnt2* and *Wnt9b* and only EC expression of *Wnt2* and *Wnt9b* appears to be evolutionarily conserved.

**Figure 1.**
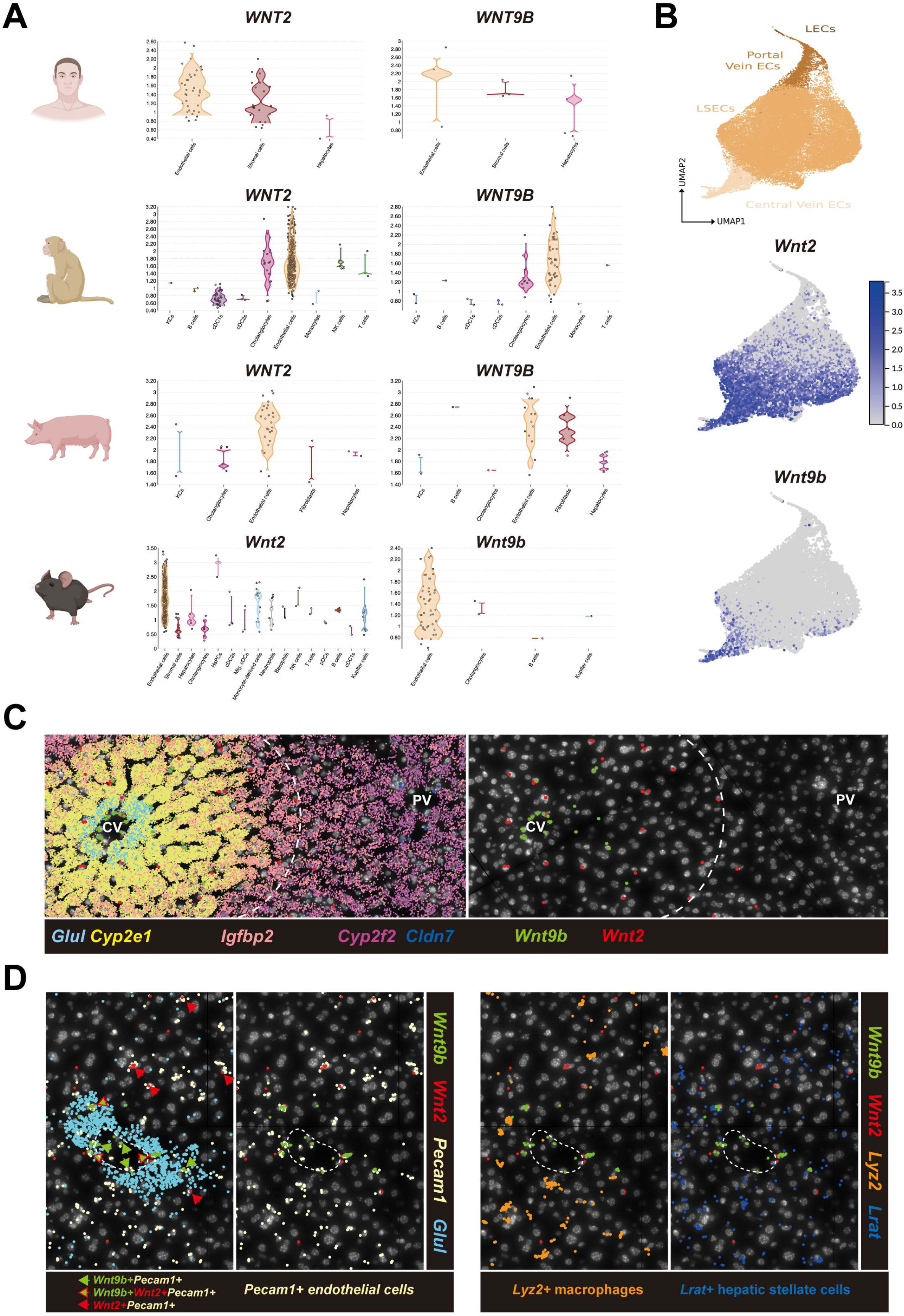
Endothelial cell expression of *Wnt2* and *Wnt9b* is evolutionarily conserved. (A) Violin plots showing expression levels of *WNT2* and *WNT9B* in human, monkey (male Cynomolgus macaques), and pig (female piglets) livers, and *Wnt2* and *Wnt9b* in mice (C57BL/6) livers among different hepatic cell types, with highest expression evident in endothelial cells across species. (Cartoons were created with BioRender.com) (B) Feature plots showing expression of *Wnt2* and *Wnt9b* among all hepatic ECs in mice. While *Wnt9b* is almost exclusively expressed in central vein endothelial cells, *Wnt2* is expressed more widely in both central vein endothelial cells as well as in liver sinusoidal endothelial cells (LSECs) towards zone 2 and zone 3. (C) Molecular Cartography showing expression in hepatocytes of *Glul and Cyp2e1* (Zone 3), *Igfbp2* (zone 2), *Cyp2f2* (zone 1) and *Cldn7* (cholangiocytes), along with *Wnt2* and *Wnt9b*, which are pericentrally zonated. (CV: central vein; PV: portal vein) (D) Molecular Cartography of *Wnt2* and *Wnt9b* along with markers of specific cell types showing *Pecam1* + ECs, but not *Lyz2+* macrophages or *Lrat+* hepatic stellate cells are the main source of *Wnt2* and *Wnt9b*.

ECs are a heterogenous population across the liver lobule (Halpern et al., 2018; Su et al., 2021). By using Uniform Manifold Approximation and Projection for Dimension Reduction (UMAP), we observed pericentrally zonated expression of *Wnt2* and *Wnt9b* in humans (Fig. S6) and in mice (Fig. 1B). Zonal expression of *Wnt2* and *Wnt9b* was also confirmed in another murine single-nuclei RNA sequencing database (Fig. S7). Consistently, *Wnt9b* was expressed mainly by central venous ECs, while *Wnt2* was expressed more broadly by both central venous ECs and hepatic sinusoidal ECs (Fig. 1B, Fig. S6).

To further validate these observations, we applied Molecular Cartography™ (Resolve Biosciences) that allows 100-plex spatial mRNA analysis on wildtype C57BL/6 mouse liver. Genes encoding for various components of the Wnt pathway and genes that are known to be zonated, were spatially resolved by Molecular Cartography™ (Table S3). We were able to identify central-portal zonation of hepatocytes using location of known zonated genes (pericentral: *Glul, Cyp2e1*; midzonal: *Igfbp2*; periportal: *Cyp2f2*) and also identify cholangiocytes based on *Cldn7* gene expression (Fig. 1C). Out of the 19 Wnts, only *Wnt2* and *Wnt9b* were pericentrally zonated (Fig. 1C). To confirm the cellular source of *Wnt2* and *Wnt9b*, we mapped gene expression of cell-specific markers including *Pecam1* for ECs, *Lyz2* for macrophages and *Lrat* for hepatic stellate cells (HSCs). *Wnt2* and *Wnt9b* predominantly colocalized with *Pecam1*, while some overlaps were also evident with *Lyz2* and *Lrat*, further confirming ECs to be the major *Wnt2* and *Wnt9b* expressing cells in zone 3 of the murine liver (Fig. 1D, Fig. S8).

Altogether, these data suggest spatially confined expression of *Wnt2* and *Wnt9b* in ECs, which along with *Rspo3* from ECs (Rocha et al., 2015) and HSCs (Dobie et al., 2019), might be instructing pericentral Wnt/β-catenin activity to contribute to metabolic LZ.

### Hepatic endothelial cell deletion of *Wnt2* and *Wnt9b* in mice

Knowing the conserved pericentral expression of *Wnt2* and *Wnt9b* in the ECs, we next sought to understand their roles via genetic elimination. *Wnt2*^flox/flox^ mice with loxP sites flanking exon 2 of the murine *Wnt2* gene were generated using CRISPR/Cas9 as discussed in methods (Fig. S9A). *Wnt9b*^flox/flox^ mice and Lyve1-Cre mice were obtained from Jackson laboratories and have been reported previously (Carroll et al., 2005; Preziosi et al., 2018). Rosa-stop^flox/flox^-EYFP mice were bred to these strains to fate-trace the expression and activity of Cre-recombinase transgene. After strategic breeding as discussed in methods, we successfully generated endothelial *Wnt2* knockout (EC-Wnt2-KO), EC-Wnt9b-KO and EC-Wnt2-9b-DKO mice, that were identified by simultaneous presence of Cre, floxed *Wnt2*, floxed *Wnt9b* and Rosa-stop alleles in the genomic DNA PCR (Fig. S9B).

While *Lyve1* is mainly expressed by midzonal hepatic sinusoidal ECs in an adult murine liver (Ma et al., 2020), Lyve1-Cre recombines floxed alleles in both hepatic sinusoidal ECs and vascular ECs likely due to expression of *Lyve1* sometime during development in all ECs (Preziosi et al., 2018). To reconfirm this, we checked GFP expression in livers from EC-Wnt2-9b-DKO mice that reflects the activity of Cre recombinase in ECs. GFP was seen in both hepatic sinusoidal ECs and central venous ECs by immunohistochemistry (IHC) and immunofluorescence (Fig. S9C, S9D). Therefore, Lyve1-Cre recombines and hence eliminates floxed alleles from both hepatic sinusoidal ECs and central venous ECs.

To examine the impact of *Wnt2, Wnt9b* or combined deletion from hepatic ECs, we assessed serum of mice from each genotype for transaminases and examined hepatic histology. Liver transaminases from all genotypes were insignificantly different from the controls supporting lack of spontaneous liver injury in any of these mice (Fig. S9E), which was also noted by lack of any histological anomaly (not shown). Liver weight to body weight ratio (LW/BW) was around 22% and significantly lower in the EC-Wnt2-KO and EC-Wnt9b-KO mice when compared to the controls, and was even lower in EC-Wnt2-9b-DKO mice, similar to what was reported for previous models of hepatic genetic elimination of various components of the Wnt/β-catenin pathway (Preziosi et al., 2018; Tan et al., 2006; Yang et al., 2014) (Fig. S9F).

### EC-derived Wnt2 and Wnt9b control expression of key □-catenin target genes in baseline liver

Since loss of β-catenin or LRP5-6 from hepatocytes or Wls from ECs, all resulted in loss of expression of prototypical Wnt/β-catenin targets in pericentral zone (Preziosi et al., 2018; Tan et al., 2006; Yang et al., 2014), we next examined EC-Wnt2-KO, EC-Wnt9b-KO and EC-Wnt2-9b-DKO mice, for similar perturbations. In males, by IHC and by western blot (WB), single deletion of *Wnt2* or *Wnt9b* led to partial but consistent loss of pericentral expression of GS, CYP2E1, and CYP1A2 (Fig. 2A, B). Minimal difference in the protein expression of these pericentral genes was evident in females (Fig. 2A, B). Interestingly, both male and female EC-Wnt2-9b-DKO mice exhibited complete loss of pericentral GS expression and had minimal residual levels of CYP2E1 and CYP1A2 (Fig. 2A). In fact, EC-Wnt2-9b-DKO phenocopied EC-Wls-KO mice in which secretion of all Wnt proteins from ECs was abolished, proving that among the 19 Wnts, *Wnt2* and *Wnt9b* are sufficient for maintaining baseline expression of key pericentral hepatocyte targets of the Wnt/β-catenin pathway and their dual deletion could not be compensated by other mechanisms (Fig. 2B).

**Figure 2.**
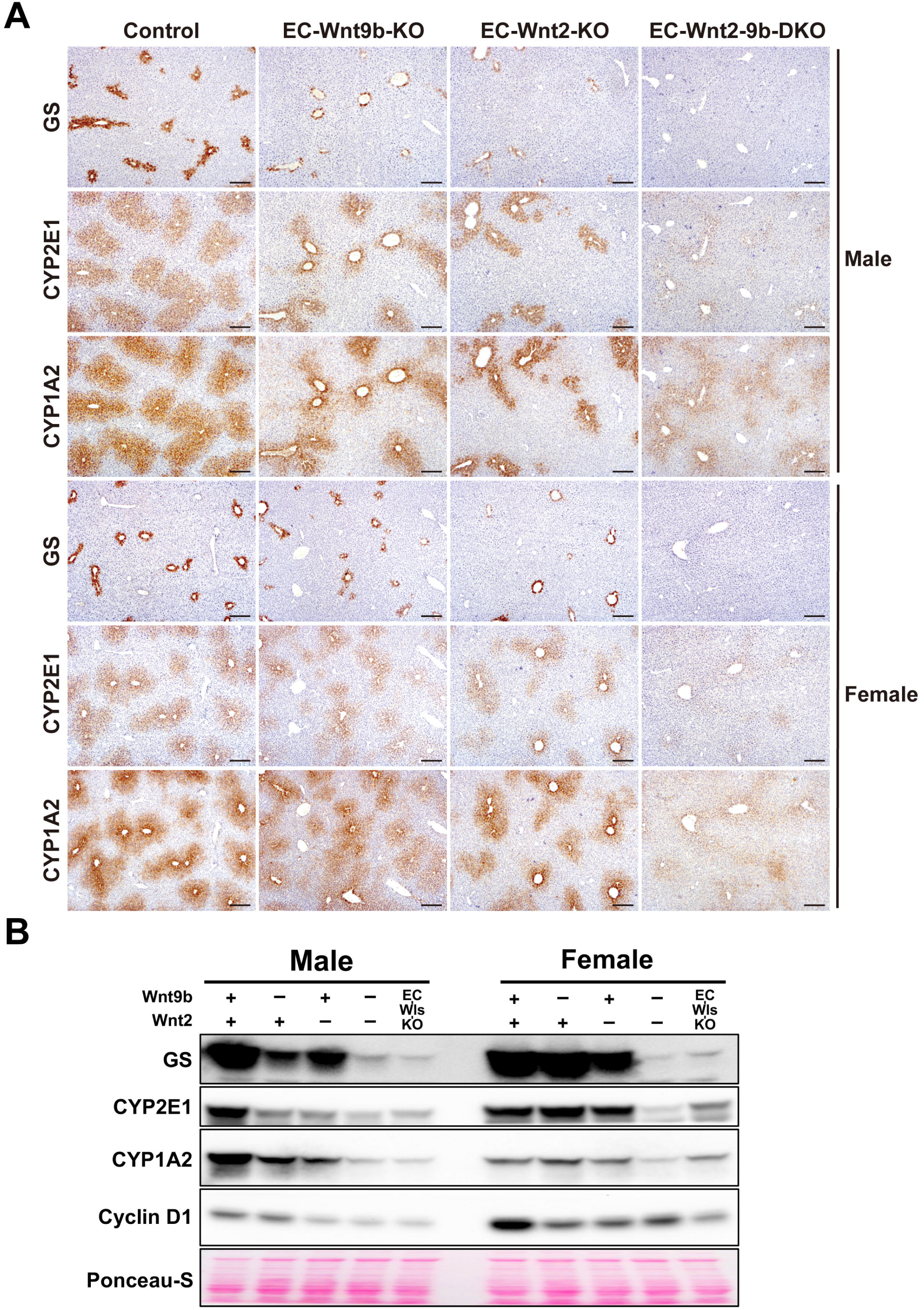
Endothelial cell-derived Wnt2 and Wnt9b control expression of key β-catenin target genes in the baseline liver. (A) Immunohistochemistry showing zonation of representative pericentral enzyme GS, CYP2E1, and CYP1A2 in representative male and female mice from age and sex-matched littermate controls (Control), EC-Wnt9b-KO, EC-Wnt2-KO and EC-Wnt2-9b-DKO. (Scale bars: 200 μm) (B) Representative WB from whole liver lysate of various genotypes showing total levels of β-catenin target genes to be modestly decreased in single EC knockouts of *Wnt2* and *Wnt9*, but more profoundly decreased in double KOs (DKOs).

Cyclin D1 is a β-catenin target gene that is mainly expressed in hepatocytes in the midzone of a quiescent liver. Like in various models of Wnt pathway perturbation in the liver (Preziosi et al., 2018; Tan et al., 2006; Yang et al., 2014), Cyclin D1 was decreased in both single and double KO mice in both males and females, but more profoundly in males and in DKOs (Fig. 2B).

### Single-cell spatial transcriptomic profiling reveals a critical role of *Wnt2* and *Wnt9b* in dynamic control of metabolic zonation

To evaluate zonation changes more comprehensively at a single-cell level, we applied Molecular Cartography™ to study the single-cell spatial expression of 100 genes (Table S3) as described above. Two pipelines were used for the analysis of the control and EC-Wnt2-9b-DKO mice (Fig. S10). First, to obtain the genetic signature at single-cell level, we used QuPath to outline single hepatocyte and obtained the expression of genes per cell. Comparable numbers of cells were obtained from control and EC-Wnt2-9b-DKO livers (Fig. 3A). Next, using expression of 16-zonated genes, a UMAP was generated which identified 6 distinct hepatocyte populations (Fig. S10). Cluster 1 and 2 represented hepatocytes with expression of pericentral genes while clusters 4, 5, and 6 represented cells expressing periportal genes (Fig. 3A). Intriguingly, only few cells exhibited pericentral gene signature in EC-Wnt2-9b-DKO and we observed a dramatic enrichment of cells in cluster 4, 5, and 6, indicating an overall shift of gene expression from pericentral to periportal in the livers with absent *Wnt2* and *Wnt9b* in ECs.

**Figure 3.**
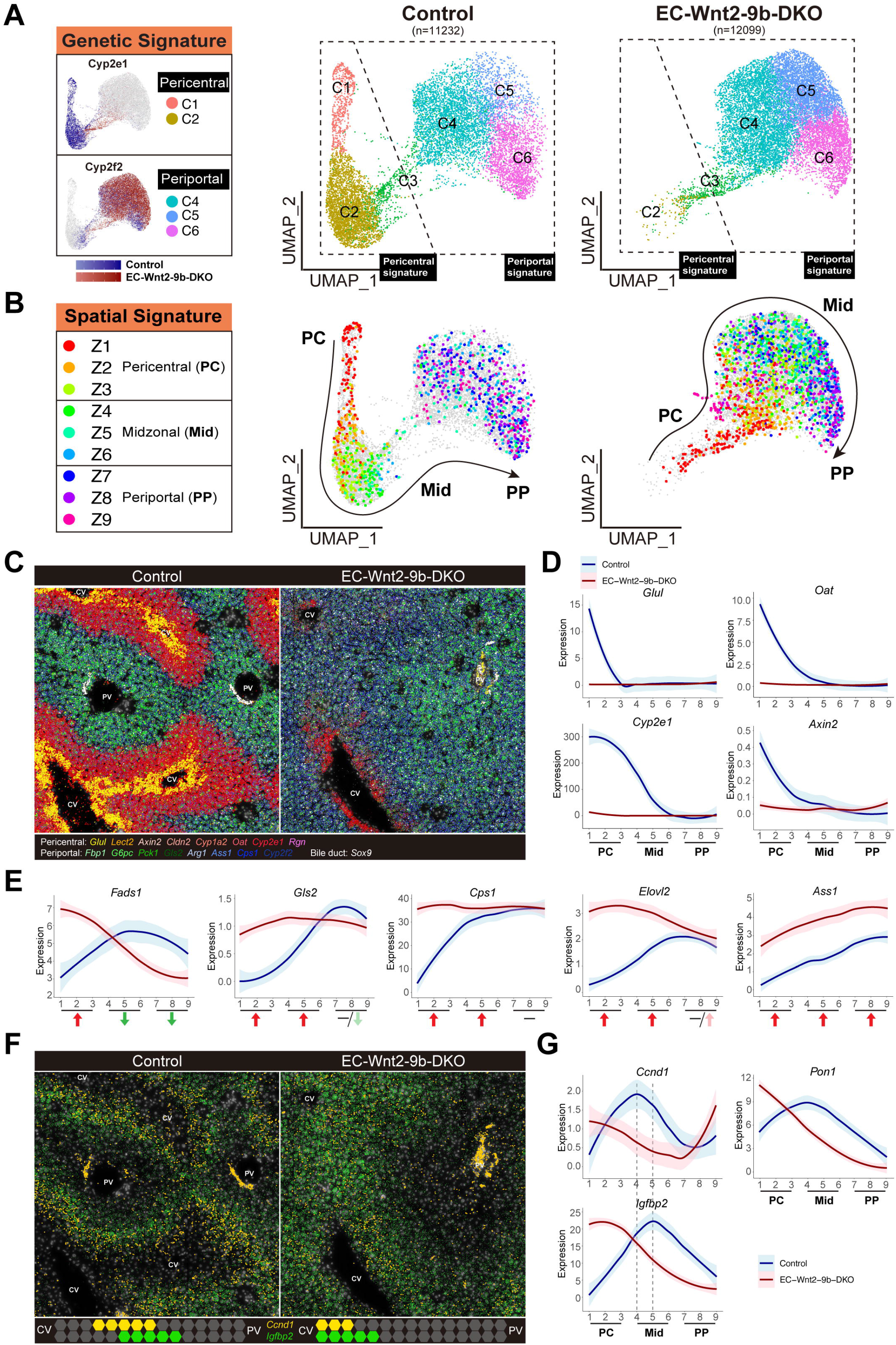
Single-cell spatial transcriptomics profiling by Molecular Cartography reveals dynamic zonation changes in EC-Wnt2-9b-DKO mice. (A) UMAP plots of single-cell transcriptomic analysis of Molecular Cartography data showing genetic clusters of hepatocytes from control and EC-Wnt2-9b-DKO livers based on expression of 16-zonated genes. Seurat clustering identifies C1-C6 clusters with C1 and C2 clusters representing hepatocytes expressing pericentral (zone 3) genes, C3 representing hepatocytes expressing midzonal genes and C4-C6 representing hepatocytes expressing periportal genes. An enrichment of C4-C6 hepatocytes is observed in the EC-Wnt2-9b-DKO at the expense of C1 and C2 clusters. (B) UMAP plots using spatial location of cells in Molecular Cartography analysis based on 9 arbitrary but equivalent zones manually drawn between portal vein and central vein shows various clusters recognized in (A) to represent zonal location validating the accuracy of this technique in addressing single-cell spatial transcriptomics. (C) Molecular Cartography comparing spatial gene expression of *Glul, Lect2, Axin2, Cldn2, Cyp1a2, Oat, Cyp2e1 and Rgn* (Zone 3), and *Fbp1, G6pc, Pck1, Gls2, Arg1, Ass1, Cps1 and Cyp2f2* (zone 1) and *Sox9* (mostly in cholangiocytes) in control and EC-Wnt2-9b-DKO liver showing loss of zone 3 genes and overall “periportalization” of the DKO liver. (CV: central vein; PV: portal vein) (D) Line plots showing impaired pericentral expression of Wnt target genes (shown here are *Glul, Oat, Cyp2e1* and *Axin2*) in DKOs as compared to control. (PC: pericentral; Mid: midzonal; PP: periportal) (E) Line plots showing dynamic changes in the expression of periportal genes (shown here are *Fads1, Gls2, Cps1, Elovl2 and Ass1*) in DKOs as compared to control. (Red arrow: upregulation; green arrow: downregulation; negative: no change. Color stands for the extent of up-or downregulation.) (F) Molecular Cartography depicting altered spatial expression of *Ccnd1* (yellow) and *Igfbp2* (green) in EC-Wnt2-9b-DKO versus control liver. While the expression of both genes is midzonal with some overlap, both genes are located pericentrally in DKOs with *Ccnd1* being more restricted to hepatocytes next to the central vein. (CV: central vein; PV: portal vein) (G) Line plots of three representative midzonal genes (*Ccnd1, Pon1* and *Igfbp2*) showing altered location to pericentral hepatocytes in DKOs as compared to control. The hue around line plots represents SEM.

Next, to obtain the spatial signature, we divided the liver lobule evenly into 9 zones (pericentral to periportal: Z1 to Z9) using landmark genes (Fig. S10). Gene expression density (gene counts per area) were quantified within each zone and averaged across the defined pericentral-to-periportal regions thus allowing us to compare gene expression across the lobule using line plots (Fig. S10). To combine the genetic signature and spatial signature, we then identified hepatocytes located within the 9 zones based on their position (X-, Y-axis) on the slides and applied this information back to the UMAP to track their localization (Fig. S10). As expected, in control mice, pericentral cells were mainly in cluster 1 and 2. Periportal cells were mainly in cluster 4, 5, and 6. However, in EC-Wnt2-9b-DKO mice, most pericentral and all midzonal and periportal cells were in cluster 4, 5, and 6, exhibiting periportal genetic signature (Fig. 3B).

The “periportalization” of the liver lobule in DKO mice was visualized by molecular cartography images of pericentral Wnt targets (*Glul, Lect2, Axin2, CIdn2, Cyp1a2, Oat, Cyp2e1, Rgn*) and periportal enzymes (*Fbp1, G6pc, Pck1, Gls2, Arg1, Ass1, Cps1, Cyp2f2*) (Fig 3C). We observed most pericentral Wnt/β-catenin target genes in hepatocytes to be dramatically downregulated or absent in EC-Wnt2-9b-DKO livers including *Glul, Oat, Cyp2e1, Axin2, Cldn2, Cyp1a2, Cyp7a1, Lect2, Lgr5, Prodh, Rgn, Rnf43, and Tbx3* (Fig 3D and Fig. S11A) supporting these to be β-catenin-TCF4 targets and under a direct control of paracrine WNT2 and WNT9B ligands from the neighboring ECs. Intriguingly, not all pericentral genes in hepatocytes were affected and paradoxically some genes like *Alad, C6, Cpox, Cyp27a1, Gstm1, and Lrp5* were upregulated (Fig. S11B) suggesting their expression to be controlled by other pericentral regulators or may represent an indirect response as part of compensation to the disruption of the Wnt/β-catenin pathway.

More interestingly, we observed *de novo* ectopic expression of most periportal genes in zone 3 either with concomitant decrease in their periportal gene expression (*Fads1, C8b, Hsd17b13, Igf1, Pck1, and Pigr*) (Fig 3E and Fig S11C), or without any change in their normal baseline periportal expression (*Gls2, Cps1, Elovl2, Arg1, Cpt2, Cyp8b1, Lrp6, Ndufb10, Uqcrh, and Vtn*) (Fig 3E and Fig S11D), or with concurrent increase in periportal gene expression (*Ass1, Atp5a1, Cyp2f2, Fbp1, Igfals, and Sox9*) (Fig 3E and Fig S11E).

Zone 2 hepatocyte proliferation has been shown to be at least in part driven by the IGFBP2-mTOR-CCND1 axis (Wei et al., 2021). Indeed, *Igfbp2* and *Ccnd1* were both expressed in midzone in the control liver, with highest expression of *Ccnd1* in Z4 and highest expression of *Igfbp2* in Z5 (Fig 3G). In EC-Wnt2-9b-DKO mice, *Ccnd1* level was overall decreased (average fold change=0.84, p=1.00654E-55), while *Igfbp2* was marginally increased (average fold change=1.09, p=1.72255E-10). Importantly, there was decrease of *Ccnd1* in midzone but an increase in pericentral hepatocytes along with an increase in *Igfbp2* (Fig 3F, 3G). The loss of *Ccnd1* from midzone in DKOs suggests its expression is also normally controlled by both pericentral *Wnt2* and *Wnt9b* from ECs and from midzonal *Igfbp2*, and the latter may become the dominant regulator in the absence of Wnts. High expression of *Ccnd1* in periportal cells is a technical artifact due to inadvertent inclusion of *Ccnd1*-positive biliary cells during cell outlining by the QuPath. Midzonal expression of *Pon1* showed a similar shift (Fig. 3G).

Collectively, changes in the expression of multiple zonated genes in hepatocytes in all zones upon elimination of *Wnt2* and *Wnt9b* that are normally expressed in zone 3 ECs underscores the overall dynamic nature of metabolic LZ. Further, LZ appears to not only be an end result of active transcription of genes but also of proactive repression of genes in the same cells.

### EC-derived Wnt2 and Wnt9b contribute to normal LR after PH

The Wnt/β-catenin pathway has also been shown to play an important role in regulating hepatocyte proliferation after PH. Loss of β-catenin or LRP5-6 from hepatocytes or Wls from ECs, all resulted in decreased Cyclin D1 expression in hepatocytes leading to notably lower numbers of hepatocytes in S-phase of cell cycle and decreased proliferation at 40 hours, which eventually recovered at 72 hours (Preziosi et al., 2018; Tan et al., 2006; Yang et al., 2014). To investigate the role of *Wnt2* and *Wnt9b* from ECs in this process, we subjected male and female EC-Wnt2-KO, EC-Wnt9b-KO and EC-Wnt2-9b-DKO mice to PH as described in methods. Male EC-Wnt2-KO and EC-Wnt9b-KO mice exhibited a profound decrease in Cyclin D1 levels (Fig. 4A). The decrease was less conspicuous in females likely due to basally higher Cyclin D1 levels (Fig. 2B, 4A). Importantly, like β-catenin-LKO, LRP5-6-LDKO, and EC-Wls-KO, Cyclin D1 was barely detectable in the EC-Wnt2-9b-DKO mice in both genders suggesting the two Wnts to be collectively essential for normal Cyclin D1 upregulation during LR (Fig. 4A).

**Figure 4.**
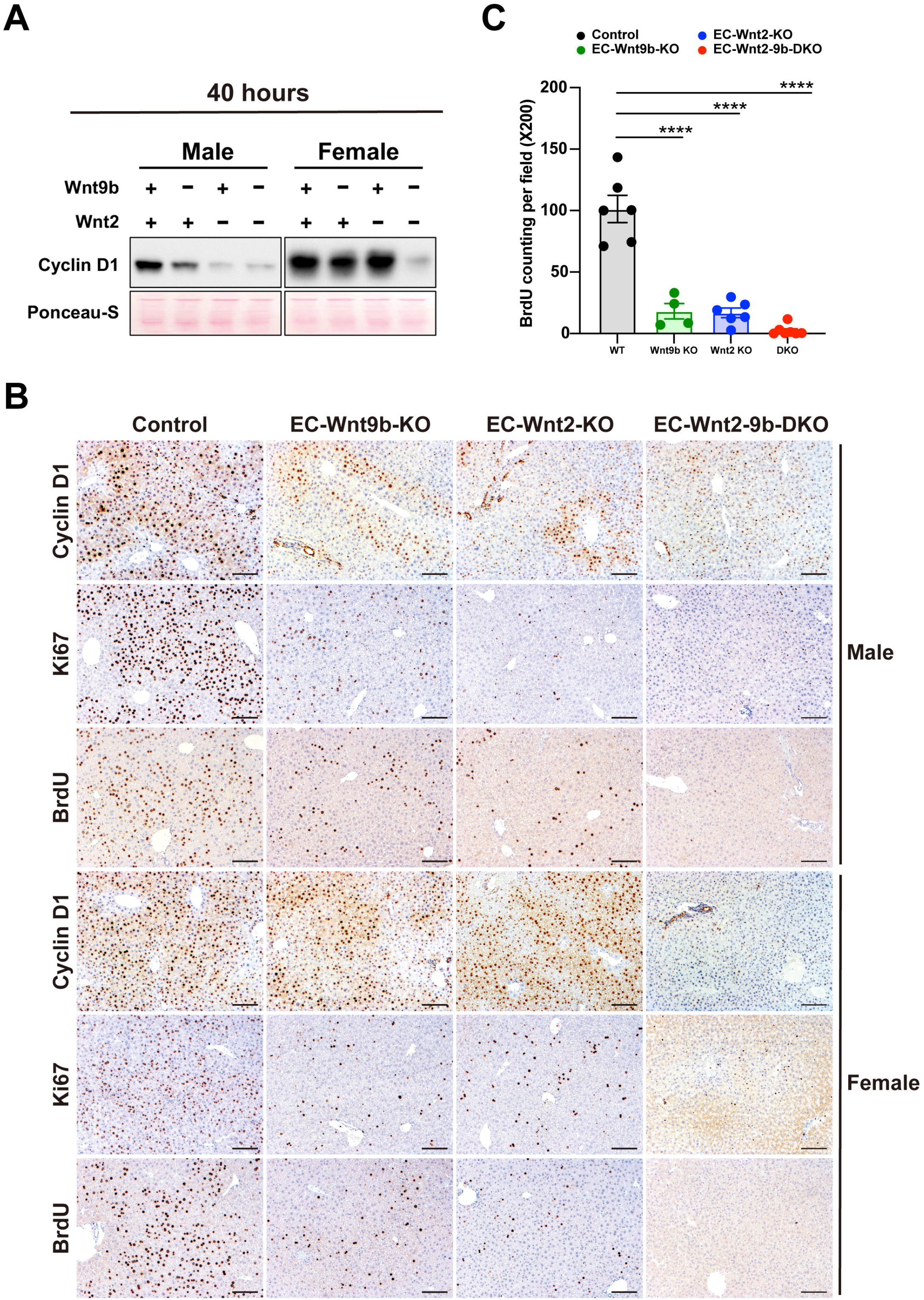
Hepatic endothelial cell-derived Wnt2 and Wnt9b contribute to normal liver regeneration after partial hepatectomy. (A) A representative WB showing notably lower Cyclin D1 level at 40h post-PH especially in males in single KO of endothelial cell Wnt2 and Wnt9b but more profound decrease is evident in DKOs in both sexes. (B) IHC for Cyclin D1 shows decreased staining at 40h post-PH in male EC-Wnt2-KO, EC-Wnt9b-KO, and EC-Wnt2-9b-DKO mice. Staining for markers of proliferation like Ki67 and BrdU was concurrently decreased in EC-Wnt2-KO and EC-Wnt9b-KO and almost absent in EC-Wnt2-9b-DKO. In females, Cyclin D1 was not changed at 40h in single KO but was notably decreases in DKOs. Despite no difference in Cyclin D1, both Ki67 and BrdU were decreased in single and almost absent in DKOs showing impairment of proliferation. (Scale bars: 100 μm) (C) Quantification of BrdU positive hepatocytes per field (200X) at 40h. Both males and females were included. (The bars represent means ± SEM. ****P < 0.0001)

We next assessed the localization of Cyclin D1 positive hepatocytes by IHC. In controls, periportal and midzonal hepatocytes were Cyclin D1 positive, while one to two layers of hepatocytes around the central vein remained negative (Fig. 4B). In single KOs, Cyclin D1 positive hepatocytes were concentrically visible around the central vein but continued to spare 1-2 layers of hepatocytes immediately around the vessel (Fig. 4B). In EC-Wnt2-9b-DKO mice, Cyclin D1 was very weakly but more diffusely expressed. These observations suggest the concentric pattern of Cyclin D1 in the EC-Wnt2-KO mice may be due to reciprocal increase in Wnt9b in pericentral neighborhood and vice versa in the EC-Wnt9b-KO mice (Fig. 4B). But their combined loss prevented such localization of Cyclin D1. Unlike males, female single KO lacked any peculiarities in Cyclin D1 levels or localization, and only DKO mice showed a dramatic decrease in Cyclin D1 expressing hepatocytes (Fig. 4B). To study the consequence of Cyclin D1 decrease and to compare to the LR in controls, we next examined single and double KO regenerating livers at 40 hours for Ki67, a marker of S-phase, and BrdU, an indicator of cell proliferation, injected 5 hours to mice before euthanasia. As what was seen for Cyclin D1, proliferating hepatocytes were mainly localized around the periportal and midzonal region in the controls. Single KO mice had sparsely positive cells, and DKO mice were completely negative for Ki67 and BrdU labeling at 40 hours (Fig. 4B, 4C). These observations were consistent in both genders. Therefore, there is a notable deficit in LR in EC-Wnt2-KO and EC-Wnt9b-KO mice, which is even worse in EC-Wnt2-9b-DKO mice phenocopying β-catenin-LKO, LRP5-6-LDKO, and EC-Wls-KO mice (Preziosi et al., 2018; Tan et al., 2006; Yang et al., 2014).

### FL6.13 rescues metabolic LZ and delayed LR in Wnt-deficient mice

Since FL6.13, the tailored tetravalent antibody has recently been shown to engage FZD-LRP6 receptor in the absence of natural Wnt ligands, we next investigated if it could activate β-catenin signaling and rescue both the pericentral gene expression or metabolic LZ as well as LR after PH in the EC-Wnt2-9b-DKO and in EC-Wls-KO mice (Preziosi et al., 2018). Four doses of control IgG or FL6.13 were injected into 8-week-old male KO or control mice as described in methods (Fig. 5A). PH was performed on day 8 and the resected livers were processed for analysis. Regenerating livers were also harvested at 24 hours post-PH. Mice were given 1 mg/ml BrdU in drinking water throughout the study to label all proliferating cells during the process (Fig. 5A). Pretreatment of FL6.13 restored pericentral expression of GS and pericentral and midzonal expression of CYP2E1 in both models (Fig. 5B). FL6.13 was also very efficient in inducing almost pan-zonal expression of Cyclin D1 barring the most immediate 1-2 layers of pericentral hepatocytes (Fig. 6B). Concomitantly, hepatocyte proliferation as seen by enhanced BrdU incorporation was evident in both KO at baseline (Fig. 5B). At 24 hours after PH, a profound increase in hepatocyte proliferation was observed in EC-Wnt2-9b-DKO and EC-Wls-KO mice by BrdU incorporation and LW/BW recovery, which was even greater than the control animals dosed with control IgG, suggesting rescue and even shift to the left in LR kinetics with FL6.13 (Fig. 5B, C). This enhanced proliferation in response to FL6.13 in both KOs was associated with continued increase in Cyclin D1 in most hepatocytes pan-zonally.

**Figure 5.**
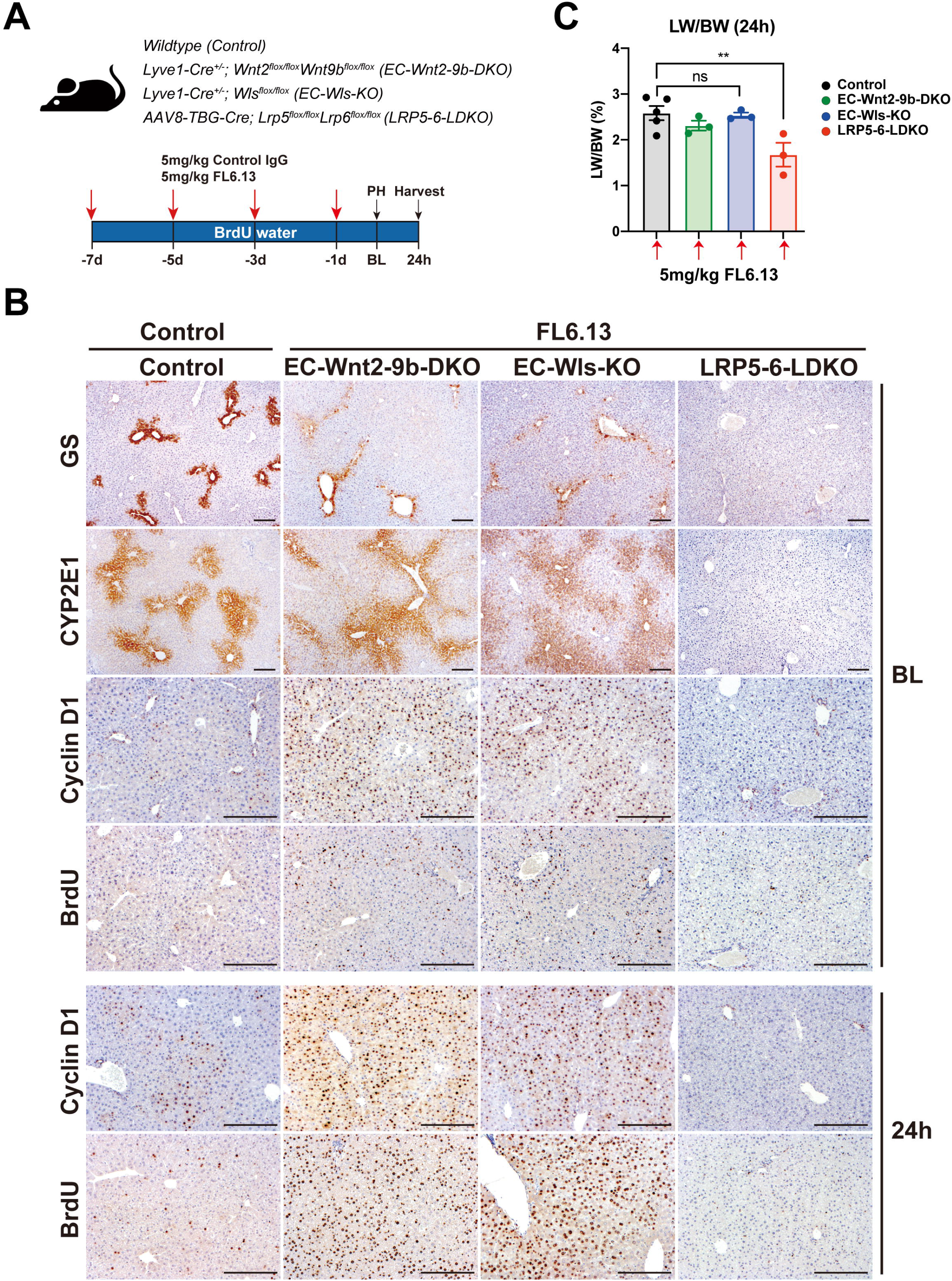
FL6.13 rescues liver zonation and liver regeneration in mice lacking Wnt secretion, specifically Wnt2 and Wnt9b from the endothelial cells, but not in mice lacking Wnt co-receptors LRP5-6 from hepatocytes. (A) Study design showing dosing schedule of Pan-Frizzled agonist FL6.13 or isotype control IgG administration to various mouse groups. (B) Representative IHC for β-catenin pericentral targets GS and CYP2E1, midzonal target Cyclin D1, and indicator of cell proliferation BrdU, in baseline livers after four treatments of control mice with control IgG and four treatments of EC-Wnt2-9b-DKO, EC-Wls-KO, and LRP5-6-LDKO mice with FL6.13. Reappearance of these markers to almost control levels is seen in EC-Wnt2-9b-DKO and EC-Wls-KO mice but not in LRP5-6-LDKO mice. (Scale bars: 200 μm) (C) Bar graph for liver weight to body weight ratio (LW/BW +/- SEM) at 24h post-PH shows comparable liver restoration in FL6.13-treated controls, EC-Wnt2-9b-DKO and EC-Wls-KO mice, but not LRP5-6-LDKO mice. (ns = not significant, **P < 0.01)

**Figure 6.**
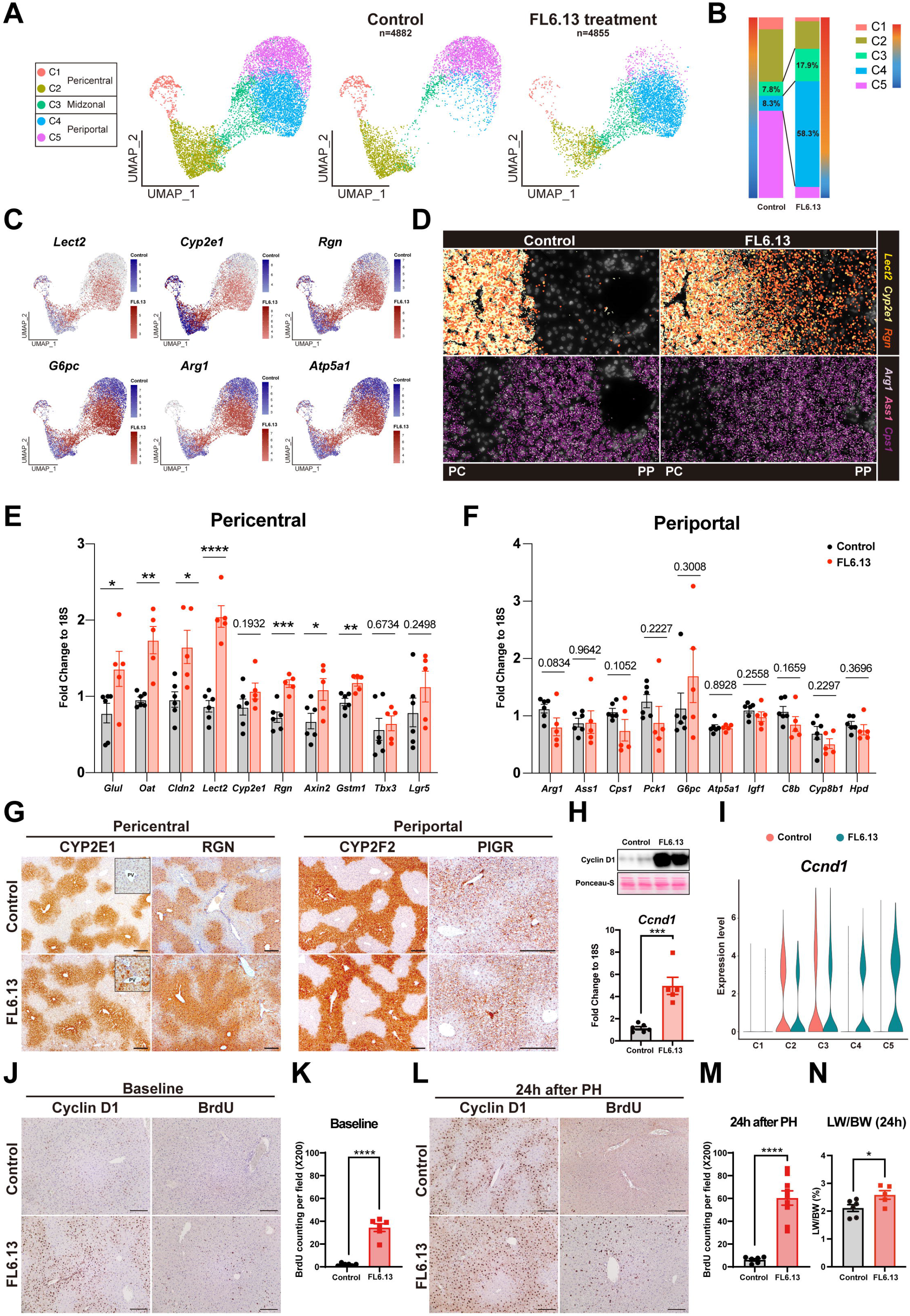
Single-cell spatial transcriptomics profiling of FL6.13-treated control mice reveals dynamic changes in zonation and promotes regeneration after partial hepatectomy. (A) UMAP plots of single-cell transcriptomic analysis of hepatocytes by Molecular Cartography showing gain of specific clusters in FL6.13-treated livers. (B) Stacked bar chart depicting distribution of various hepatocyte clusters showing specific gain of C3 and C4 after FL6.13 treatment (right bar) as compared to controls (left bar). (C) Feature plots from Molecular Cartography analysis of specific representative genes showing ectopic expression of pericentral genes (*Lect2, Cyp2e1, Rgn*) in midzonal and periportal cluster after FL6.13 treatment (red) as compared to controls (blue), while periportal genes (*G6pc, Arg1, Atp5a1*) continued to be expressed in these clusters and hence remained unchanged. (D) Molecular Cartography visualization of tissue section of the same genes as indicated in (C) showing ectopic expression of pericentral genes in midzonal and periportal regions while periportal genes were unchanged after FL6.13 treatment. (PC: pericentral; PP: periportal) (E) qPCR using bulk mRNA from livers of controls or FL6.13-treated mice shows significant increase in the expression of several pericentral Wnt target genes by FL6.13. (*P < 0.05, **P < 0.01, ***P < 0.001, ****P < 0.0001) (F) qPCR using bulk mRNA from livers of controls or FL6.13-treated mice shows no statistically significant effect on expression of several known periportal genes by FL6.13. (Corresponding P values are indicated) (G) Representative IHC verify ectopic localization of CYP2E1 and RGN proteins in midzonal and periportal regions after FL6.13 treatment, while CYP2F2 and PIGR protein locations remained unchanged. (PV: portal vein. Scale bars: 200 μm) (H) Representative qPCR using whole liver RNA and WB using whole liver lysate shows dramatic increase in *Ccnd1* expression and Cyclin D1 protein after FL6.13 treatment as compared to controls. (I) Violin plot of Seurat clustering of the Molecular Cartography from (A) shows appearance of periportal *Ccnd1* expression (C4 and C5 clusters; green) while its expression in C2 and C3 clusters where it is normally expressed in controls (red), remains unchanged after FL6.13 treatment (green). (J) Representative IHC for Cyclin D1 and BrdU showing FL6.13 induces periportal hepatocyte G1 to S phase transition and hepatocyte proliferation, respectively as compared to controls. (Scale bars: 100 μm) (K) Quantification of BrdU positive hepatocytes (+/-SEM) in (J) shows FL6.13 induces significant increase in hepatocyte proliferation at baseline as compared to control IgG. (****P < 0.0001) (L) Representative IHC showing FL6.13 accelerates liver regeneration at 24h post-PH by promoting number of Cyclin D1-positive periportal and midzonal hepatocytes and concurrently increasing BrdU-positive proliferating periportal and midzonal hepatocytes, as compared to control IgG treatment. (Scale bars: 100 μm) (M) Quantification of BrdU positive hepatocytes (+/-SEM) in (L) shows FL6.13 induces significant increase in hepatocyte proliferation at 24 hours post hepatectomy as compared to control IgG treatment. (****P<0.0001) (N) LW/BW (+/-SEM) at 24h post-PH depicts an advantage of liver mass restoration in the FL6.13 versus control IgG-treated group. (*P < 0.05)

To address specificity of the response by FL6.13, we next treated LRP5-6-LDKO mice lacking Wnt-co-receptors in hepatocytes (Yang et al., 2014). These mice have been previously shown to also phenocopy β-catenin-LKO in both lacking pericentral Wnt/β-catenin targets and delayed LR. When LRP5-6-LDKO were treated similarly with FL6.13 (Fig. 5A), there was no change in pericentral gene expression of GS and CYP2E1 which continued to be absent in these mice (Fig. 5B). Similarly, FL6.13 was unable to restore either Cyclin D1 expression nor BrdU incorporation either in baseline or 24 hours post-PH livers (Fig. 5B). This was also reflected by a deficient LW/BW at 24 hours post-PH in the LRP5-6-LDKO mice as compared to controls or FL6.13-treated EC-Wnt2-9b-KO mice (Fig. 5C). This study shows the requirement of intact Wnt co-receptors for FL6.13 to stimulate the Wnt/β-catenin signaling *in vivo*.

### FL6.13 induces pericentral gene expression at baseline

To characterize Wnt/β-catenin activation by FL6.13 in greater depth, we tested the effect of four doses of FL6.13, given every 48 hours, on the wildtype mice as described in the methods. Twenty-four hours after the last injection, the mice were euthanized, and livers were processed for Molecular Cartography™ using the same set of probes (Table S3). Two pipelines were applied for analyses (Fig. S12). Zonal distribution of hepatocytes was identified based on differentially expressed genes (Fig. S13) and confirmed by tracing back the localization and zonation of hepatocytes on the slides (Fig. S12). The genetic signature and spatial signature overlapped quite well on the UMAP for the current analysis unlike the EC-Wnt2-9b-DKO mice that lacked Wnt/β-catenin target genes in zone 3 hepatocytes precluding overlap (Fig. S12). Five different clusters were identified, which enabled reconstruction of the metabolic zones (pericentral: cluster 1 and 2; midzonal: cluster 3; periportal: cluster 4 and 5) (Fig. 6A). A notable increase of the proportion of C3 and C4 was noted after FL6.13 treatment (Fig. 6B), which was marked by the ectopic expression of pericentral Wnt target genes such as *Lect2, Cyp2e1, Rgn, Cyp1a2, Gstm1*, and *Cldn2* (Fig. 6C and Fig. S14A). Interestingly, some pericentral genes were not induced by FL6.13 using spatial single-cell analysis, including transcription factor *Tbx3* (Renard et al., 2007), heme synthesis enzymes *Alad* and *Cpox* (Braeuning and Schwarz, 2010), bile acid synthesis enzyme *Cyp27a1* (Halpern et al., 2017), and complement pathway gene *C6* (Halpern et al., 2017) (Fig. S14B). This indicated that FL6.13 selectively activates some but not all pericentral genes. Unlike de-repression of periportal genes in the EC-Wnt2-9b-DKO mice, there was no concomitant decrease in the expression of periportal genes including complement pathway genes (*C8b, Vtn*), Fc receptor (*Pigr*), lipid metabolism gene (*Hsd17b13*), oxidative phosphorylation genes (*Uqcrh, Atp5a1*), gluconeogenesis genes (*G6pc, Fbp1, Pck1*), glutamine catabolism gene (*Gls2*), urea cycle genes (*Arg, Cps1*), and hormone (*Igf1*) (Fig. 6C and Fig. S15A). Representative images from the Molecular Cartography™ showed that FL6.13 induced an expansion of pericentral markers (*Lect2, Cyp2e1, Rgn*) to both midzonal and periportal regions, while the expression of periportal markers (*Arg1, Ass1, Cps1*) remained unaltered (Fig. 6D).

We also validated some of these findings by qPCR. Indeed, most pericentral Wnt/β-catenin target genes were significantly induced by FL6.13 treatment, whereas periportal genes were not affected (Fig. 6E and 6F). Not all pericentral genes were increased by FL6.13. Levels of *Tbx3* and *Lgr5* were unchanged after FL6.13 treatment (Fig. 6E).

To visualize zonation changes at the protein level, we stained pericentral markers (CYP2E1 and RGN) and periportal markers (CYP2F2 and PIGR) in control and FL6.13-treated livers. Expansion and ectopic expression of CYP2E1 and RGN was observed at the midzone and in the periportal hepatocytes after FL6.13 treatment, while areas of CYP2F2 and PIGR immunostaining were similar as controls and in periportal hepatocytes (Fig. 6G).

Altogether, these data indicate a selective and potent effect of FL6.13 in inducing expansion of most pericentral Wnt/β-catenin target genes but not at the expense of periportal gene expression.

### FL6.13 induces *Ccnd1* gene expression, hepatocytes proliferation and promotes LR after PH

Since FL6.13 effectively promoted LR after PH in the EC-Wnt2-9b-DKO and EC-Wls-KO mice through stimulation of *Ccnd1* expression, we next queried its impact on normal baseline liver after four doses of FL6.13 given every 48 hours. A notable increase in *Ccnd1* mRNA and Cyclin D1 protein was observed after FL6.13 treatment (Fig. 6H). Interestingly, induction of Cyclin D1 was not evenly distributed across the liver lobule, but only around the periportal region (Fig. 6I, 6J), similar to observations with another Wnt agonist and RSPO1 (Janda et al., 2017; Planas-Paz et al., 2016). BrdU incorporation was significantly induced after FL6.13 treatment, indicating that periportal hepatocytes entered S-phase upon Wnt activation and Cyclin D1 expression (Fig. 6J, K). Collectively, FL6.13 could activate hepatic Wnt/β-catenin pathway, induce expansion of pericentral metabolic LZ through midzone and even periportally, without impairing expression and localization of periportal genes, and while provoking proliferation of periportal and midzone hepatocytes. These data also suggest that hepatocytes within different zones in the liver have distinct responses to exogenous Wnt stimulation.

Considering the strong effect of FL6.13 in inducing hepatocyte proliferation even at baseline, as a proof-of-concept for therapeutic intervention, we next investigated the effect of FL6.13 on LR using the PH model. Four doses of control IgG or FL6.13 were i.p. injected to 8-week-old male C57/BL6. PH was performed on day 8 and regenerating livers were harvested at 24 hours. Mice were given 1 mg/ml BrdU in drinking water to label proliferating cells throughout the process (Fig. S15B). In control IgG-treated mice, Cyclin D1 was expressed mainly at the midzone at 24 hours post-PH, and very few hepatocytes had BrdU positive nuclei (Fig. 6L). Pretreatment of FL6.13 induced a dramatic expansion of Cyclin D1 towards pericentral region and six-fold increase of periportal BrdU incorporation (Fig. 6L, 6M). Proliferative advantage led to accelerated LR and contributed to significantly greater recovery of LW/BW at 24-hour time point (Fig 6N). Therefore, these results demonstrate a potent effect of FL6.13 pretreatment in promoting LR leading to faster recovery of hepatic mass.

### Late treatment with FL6.13 promotes liver repair after acetaminophen injury by promoting Wnt/β-catenin activation and hepatocyte proliferation

To test the clinical applicability of FL6.13 to promote repair in a model of acute liver insult, we evaluated its efficacy in acetaminophen (APAP) overdose-induced hyperacute liver injury. APAP is a widely used analgesic and antipyretic, but it accounts for 46% of acute liver failure (ALF) in the United States (Lee, 2017). Patients with APAP overdose either progress to liver failure requiring transplant or may recover spontaneously. N-acetyl cysteine (NAC) is the only currently approved antidote but must be given early after APAP ingestion for its full efficacy (Ershad et al., 2022), and the probability of developing ALF continues to increase over time with delay in NAC administration.

Eight-week-old male C57/BL6 fasted for 12 hours were injected 600 mg/kg APAP followed by a single dose of control IgG or FL6.13 at 12 hours and mice were sacrificed at 48 hours post APAP injection (Fig. S16A). Surprisingly, mice treated with FL6.13 had higher mortality (Fig. S16B), accompanied with two-fold higher ALT level (Fig. S16C). Negative effect of FL6.13 was seen as patches of necrotic areas by HE (Fig. S16D). Severe gross and sinusoidal congestion was seen in the dead mice after FL6.13 treatment (Fig. S16B, D). We next evaluated hepatocyte proliferation that counteracts the injury to drive liver repair. 600 mg/kg APAP has been reported to induce severe liver injury with mild hepatocyte proliferation during the recovery process (Bhushan et al., 2014). Indeed, in the control group Cyclin D1 was weakly expressed only by the first layer of hepatocytes surrounding pericentral necrotic regions at 48 hours post-APAP. Hepatocytes were negative for Ki67 demonstrating lack of G1 to S-phase transition at this time (Fig. S16D). Interestingly, FL6.13 treatment did not induce Cyclin D1 and Ki67 at 48 hours after APAP-overdose (Fig. S16D). Comparable immune cell infiltration, important for phagocytosis of necrotic cells and subsequent liver repair (Jaeschke and Ramachandran, 2020), was evident in both control IgG and FL6.13-treated mice, shown by IHC of CD45 (Fig. S16D). CYP2E1 plays an essential role in APAP metabolism to generate N-acetyl-p-benzoquinone imine (NAPQI), a highly reactive metabolite that leads to hepatotoxicity (Yoon et al., 2016). Since FL6.13 was shown to induce pericentral and periportal CYP2E1 expression in wildtype mice (Fig. 6G), it is likely that 12 hours post 600 mg/kg APAP injection, remnant APAP in mice was inadvertently metabolized to NAPQI by FL6.13-induced CYP2E1 expression, leading to a more severe liver injury. Our data suggest that early intervention with a Wnt agonist may have unintended consequences and may worsen APAP-induced liver injury due to induction of CYP2E1.

Next, we investigated if late treatment of FL6.13 could have any therapeutic benefit through promoting regeneration especially when APAP has already been metabolized. Such opportunity represents an unmet clinical need since there are no therapies currently available if NAC fails to prevent ALF when patients arrive for clinical intervention late after APAP overdose. Single dose of control IgG or FL6.13 was administrated at 32 hours post 600 mg/kg APAP injection, and mice were sacrificed at 60 hours (Fig. 7A). Grossly, livers displayed normal color and appeared healthier in the FL6.13 group (Fig. 7B). FL6.13-treated mice showed significantly reduced serum ALT while AST trended favorably as well (Fig. 7C). Control group showed pericentral necrotic areas at 60 hours post APAP injection with increased expression of Cyclin D1 in peri-necrotic hepatocytes (Fig. 7D). Ki67 positive hepatocytes were seen surrounding necrotic region and also in hepatocytes at the periportal region (Fig. 7D). FL6.13-treated mice exhibited a significantly reduced necrotic area, which was associated with a profound increase in Cyclin D1 expressing hepatocytes pan-zonally indicating these cells are all able to enter cell cycle across the liver lobule (Fig. 7D, 7E). Many periportal hepatocytes as well as pericentral hepatocytes were Ki67 positive in the FL6.13-treated group, which was significantly more than the isotype control-treated group (Fig. 7D, F). FL6.13 treatment showed a more localized immune cell response around necrotic regions as seen by staining for CD45 (Fig. 7D). Altogether, delayed Wnt agonism promotes liver repair after acute APAP injury by inducing hepatocyte proliferation that occurs locally and periportally and may be ‘pushing’ hepatocytes towards central vein to fill the space created by the cleared necrotic debris by the immune cells. Thus, these data extend the application of FL6.13 as delayed regenerative treatment choice in APAP associated ALF and other relevant indications.

**Figure 7.**
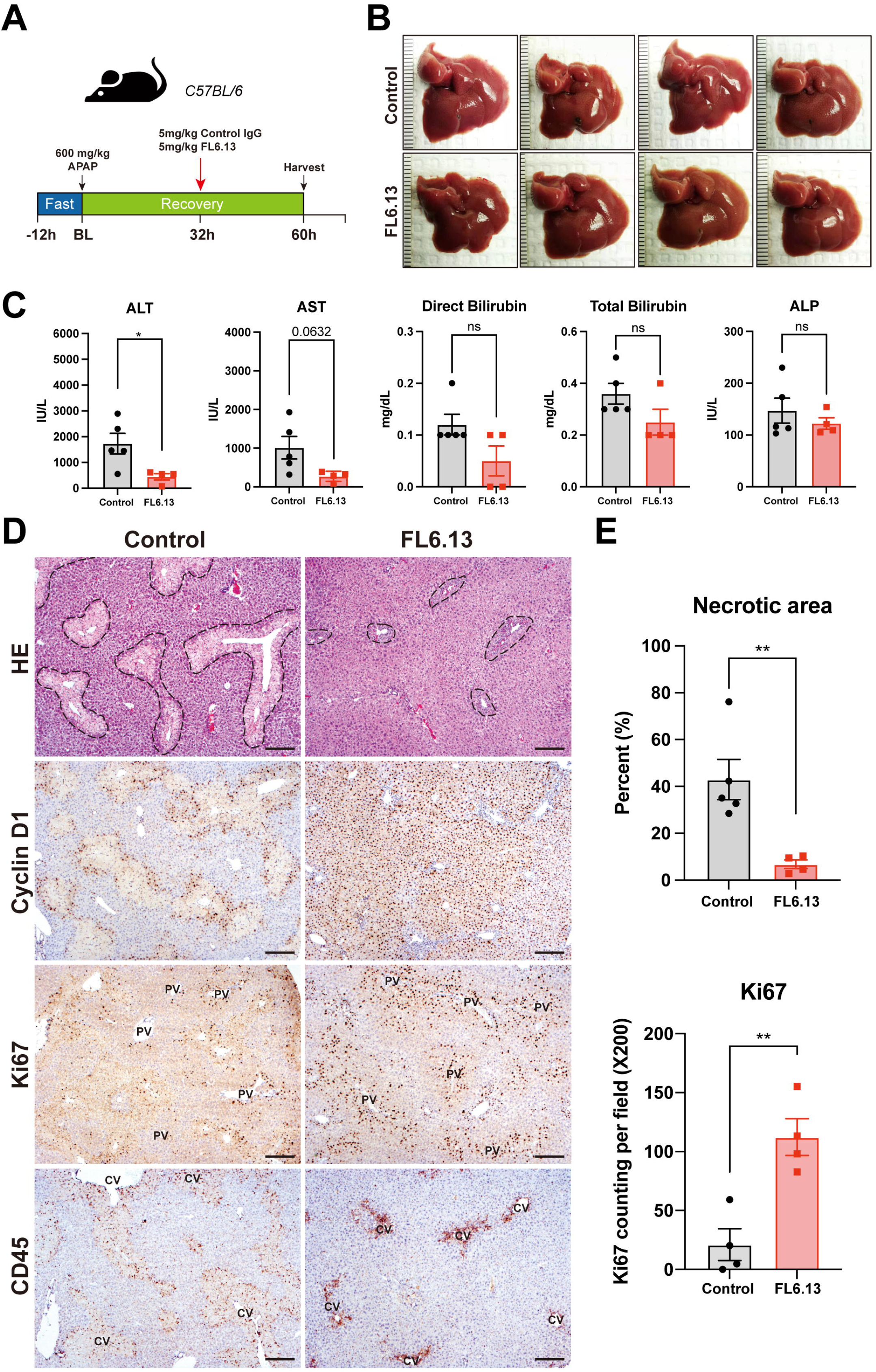
Late treatment with FL6.13 promotes liver repair after acetaminophen injury through induction of hepatocyte proliferation. (A) Study design showing administration of a single dose of 5 mg/kg pan-FZD agonist FL6.13 or isotype control IgG at 32 hours post 600 mg/kg i.p. APAP injection. Mice were sacrificed at 60 hours for analysis. (B) Gross images of livers from FL6.13 and control IgG treated groups after APAP showing decreased necrosis and congestion at 60 hours in the FL6.13 treatment group. (C) Serum ALT, AST, direct and total bilirubin, and ALP levels (+/-SEM) at 60 hours after APAP showing significantly reduced serum ALT and AST (trending favorably) in FL6.13 as compared to control IgG. (ns = not significant, *P < 0.05) (D) Representative IHC showing decreased necrotic areas by Hematoxylin and Eosin (HE) staining, pan-lobular Cyclin D1 staining, increased periportal cell proliferation by Ki67 staining, and a more localized immune cell response by CD45 staining after FL6.13 treatment when compared to control IgG. (CV: central vein; PV: portal vein. Scale bars: 200 μm) (E) Quantification of HE and Ki67 showing significant decrease in necrotic areas and increase in hepatocyte proliferation in FL6.13 group. (**P < 0.01)

## Discussion

Substantial studies place β-catenin in the center of liver pathophysiology, with its unique roles in maintaining LZ and promoting LR. In view of this, it is of importance to answer “Who regulates the regulator?”. While the observation of *Wnt2* and *Wnt9b* being present at the central venous ECs and pericentral sinusoidal ECs (Wang et al., 2015), and their upregulation in various liver injuries (Ding et al., 2010; Preziosi et al., 2018; Zhao et al., 2019) have suggested their important roles, without genetic tools, it was not possible to definitively and unequivocally demonstrate their contributions in LZ and LR. Here, we provided the genetic evidence and thus conclusively showed that ECs-derived *Wnt2* and *Wnt9b* play an additive and equivalent role in instructing metabolic zonation and in driving hepatocyte proliferation after PH. Although *Wnt2* is expressed more broadly than *Wnt9b*, their role appears to be additive, especially in males, and overall indistinguishable from each other in the current analysis. Interestingly, we observed gender differences such that females were able to compensate single loss of *Wnt2* or *Wnt9b* better than males, and the underline mechanism requires further investigation.

Although previous studies have hinted that periportal enzymes could be ectopically expressed by zone 3 hepatocytes upon β-catenin inhibition, the conclusions have been limited by unavailability of technical tools (Benhamouche et al., 2006). Until now, it has not been possible to comprehensively access gene expressions at the single-cell level in a high-throughput manner, restricted either by spatial resolution like Visium, or by the high-throughput ability like single-molecule FISH, all of which limited the in-depth understanding of the dynamic zonation. By using Molecular Cartography™, we were able to evaluate spatial expression of hundred genes on the same slide and analyze their expression changes upon genetic or pharmacological intervention at cellular resolution, thus creating single-cell spatial transcriptomic profiling. We extended these findings to cellular and spatial context, and interestingly, found that in EC-Wnt2-9b-DKO mice, many analyzed periportal genes were induced *de novo* in the zone 3 hepatocytes. Intriguingly, this occurred simultaneously with changes in their expression even in their native neighborhoods. While some genes showed concurrent increase in their native zone, others showed no change, and yet others showed concomitant decrease. This shows the process of metabolic LZ to be highly dynamic and suggests that upstream effectors and master regulators of the process are not only responsible for actively transcribing genes, while proactively repressing others. Interestingly, EC-Wnt2-9b-DKO mice showed periportalization of the liver, however, administration of the pan-FZD agonist, while it induced pericentralization of the liver by inducing zone 3 gene expression periportally, it did not result in loss of zone 1 gene expression. One possible explanation of the difference could be permanent genetic elimination versus transient effect brought about by administration of an exogenous molecule. Another possibility is the differential zonal dynamism of gene expression in zone 3 versus zone 1 such that progenitors during development spontaneously give rise to hepatocytes with default expression of periportal genes, whereas zone 3 hepatocyte identity is acquired by neighborhood signals such as from zone 3 ECs which leads to simultaneous repression of periportal genes and activation of pericentral genes. Indeed, hepatocytes derived from differentiation of liver stem cells have spontaneous periportal gene expression while pericentral gene expression needs to be forced by β-catenin activation (Colletti et al. 2009).While further studies are needed to address how zone 3 hepatocytes shift to periportal identity in the EC-Wnt2-9b-DKO mice, previous studies have also shown that HNF4α inhibits pericentral β-catenin target gene expression (Stanulović et al., 2007). In the absence of β-catenin, TCF4 associated with HNF4α could possibly bind to HNF4α responsive elements (HREs), inducing the expression of periportal genes (Berasain and Avila, 2014; Gougelet et al., 2014), and could be one plausible explanation for our observations.

Another important question that remains unanswered is what regulates *Wnt2* and *Wnt9b* expression and secretion at baseline in the ECs in zone 3, and what stimulates their immediate-early upregulation and release after PH. Likewise, why *Wnt2* and *Wnt9b* are the major ‘regenerative and homeostatic’ Wnts in the liver and not others? It is likely that there are specific upstream regulators unique to *Wnt2* and *Wnt9b* gene expression, and further studies will be essential to unravel such effectors. Currently, there exist some indications of role of Gata4 in hepatic sinusoidal endothelial *Wnt2* expression whose deletion resulted in impaired β-catenin signaling and disturbed LZ (Winkler et al., 2021). Systemic blockade of vascular receptor tyrosine kinase Tie1, but not VEGFR-2, VEGFR-3, Dll4, integrins-αV, integrin-α5, or PECAM1 has been shown to abrogate *Wnt9b* from central venous ECs within 2 hours (Inverso et al., 2021). ECs-specific *Tie1* deletion also delayed LR due to impaired induction of *Wnt2* and *Wnt9b* (Inverso et al., 2021). Notch has also been shown to regulates c-kit positive ECs to contribute to LZ and LR through *Wnt2* (Duan et al., 2022). Finally, deletion of endothelial Heart-of-glass (Heg) changed vascular 3D-patterning and disrupted LZ through impaired *Wnt2, Wnt9b*, and *Rspo3* expression (Zhu et al., 2022). Whether these various perturbations are because each of these regulates *Wnt2* and *Wnt9b* expression and/or secretion from ECs or it is due to effect of these alterations on endothelial cell function and identity leading to secondary changes in Wnts, remains unclear and requires further careful analysis.

Being a critical pathway in modulating both hepatocyte metabolism and proliferation, activating the Wnt/β-catenin to promote hepatic function and restore mass is an attractive therapeutic strategy. We utilized a novel pan-FZD agonist FL6.13, a tetravalent antibody described recently, which induces FZD-LRP6 engagement to mimic physiological Wnt activation circumventing the need of Wnts (Tao et al., 2019). Indeed, FL6.13 rescued pericentral LZ in EC-Wnt2-9b-DKO and EC-Wls-KO mice but not in the LRP5-6-LDKO mice. FL6.13 modulated hepatic metabolism leading to expansion of pericentral Wnt targets in zone 1 and zone 2 without the expense of normal gene expression in the periportal region. It was able to simultaneously induce *Ccnd1* expression, allowing hepatocytes to enter cell cycle and in this way stimulated liver repair without impacting metabolic function. This duality of a Wnt agonism to induce cell proliferation while maintaining metabolic function could have specific role in treating both acute and chronic liver insufficiency (Hu and Monga, 2021).

One specific and highly relevant unmet need is delayed treatment of patients of APAP overdose. In fact, the chances of development of hepatic failure continues to increase despite administration of NAC, the only approved therapy for APAP overdose. If given within 4-10 hours of excessive APAP ingestion, hepatoxicity could still develop in 6.1% cases given NAC. The chances of developing failure rise to 26.4% and 41%, if NAC is administered 10-24 hours or beyond 24 hours of APAP overdose, respectively (Smilkstein et al., 1988). There is no effective therapy beyond 24 hours and patients will either spontaneously recover or will progress requiring transplant. Our data provides a new treatment option and fulfills this unmet need of administering regenerative therapy to cases who are either delayed seeking medical attention or progress to failure despite NAC. The advantage of use of pan-FZD agonist like FL6.13 is that it enhances pericentral metabolic zonation while simultaneously promoting both periportal and pericentral hepatocyte proliferation. The periportal hepatocyte proliferation may be creating the push needed for hepatocytes to move pericentrally to restore microarchitecture in this area as the necrotic debris gets cleared by immune cells. However, it is important to note that premature Wnt stimulation could inadvertently induce genes like CYP2E1 and CYP1A2 to generate excessive APAP toxic metabolites and hence timing of intervention with FL6.13 like agents would need to be carefully selected. Additionally, serum APAP levels might need to be tested before administration of any Wnt agonist. However, regenerative therapy may fill an important niche in treatment of cases of ALF due to APAP and perhaps other causes.

## Author contributions

Conceptualization, S.H. and S.P.M.; Methodology, S.H., S.L., Y.B., M.P., S.S., C.C., J.M., A.B., and S.P.M.; Investigation, S.H., S.L., Y.B., M.P., S.S., C.C., J.M.; Writing, S.H. and S.P.M.; Funding Acquisition, S.P.M; Resources, L.L.B., J.J.A., S.S.S., and S.A.

## Declaration of interests

Dr. Monga is a consultant for Surrozen. Drs. Blazer, Adams, Sidhu and Angers are shareholders of AntlerA Therapeutics.

## Supplementary figure legends

**Fig. S1 Violin plots showing expression level of WNTs in normal human liver.** (Related to Figure 1)

Eighteen WNTs were detected by scRNA seq. ECs were the predominant source of *WNT2, WNT2B*, and *WNT9B*.

**Fig. S2 Violin plots showing expression level of WNTs in monkey liver.** (Related to Figure 1)

Sixteen WNTs were detected. ECs expressed high levels of *WNT2* and *WNT9B*.

**Fig. S3 Violin plots showing expression level of WNTs in pig liver.** (Related to Figure 1)

Nine WNTs were detected. ECs expressed high levels of *WNT2* and *WNT9B*.

**Fig. S4 Violin plots showing expression level of Wnts in C57/BL6 mice liver.** (Related to Figure 1)

Fifteen Wnts were detected. ECs expressed high levels of *Wnt2* and *Wnt9b*.

**Fig. S5 Violin plots showing expression level of Wnts in NAFLD mice liver.** (Related to Figure 1)

Fourteen Wnts were detected. Endothelial expression of *Wnt2* and *Wnt9b* was maintained in NAFLD mice. (Cartoons were created with BioRender.com)

**Fig. S6 Feature plots showing expression and distribution of WNTs in human liver.** (Related to Figure 1)

*WNT2* was pericentrally zonated in human ECs.

**Fig. S7 Feature plots showing expression and distribution of Wnts in C57/BL6 mice liver using single nuclei RNA sequencing.** (Related to Figure 1)

Among thirteen detected Wnts, only *Wnt2* and *Wnt9b* were pericentrally zonated by snRNA seq.

**Fig. S8 Molecular Cartography of indicated genes.** (Related to Figure 1)

Colocalization of *Wnt2* and *Wnt9b* with cell-specific markers including *Pecam1* for ECs, *Lyz2* for macrophages, *Lrat* for hepatic stellate cells, and *Ptprc* for immune cells. *Wnt2* and *Wnt9b* predominantly colocalized with *Pecam1*, while some overlap was also evident with *Lyz2, Lrat, and Ptprc.* (CV: Central vein; PV: portal vein)

**Fig. S9 Characterization of mice models.** (Related to Figure 2)

(A) Scheme depicting loxP site (Red lines) in mice *Wnt9b* and *Wnt2* gene. (Black boxes: exons; grey boxes: 5’-UTR and 3’-UTR non-coding regions; arrow heads: PCR primers for genotyping)

(B) Gel electrophoresis showing generation of mice as indicated in the figure. (C, D) IHC and immunofluorescence of EC-Wnt2-9b-DKO mice showing hepatic sinusoidal ECs and central venous ECs were positive for GFP. (CV: central vein; PV: portal vein. Scale bars for IHC: 20 μm; Scale bars for IF: 100 μm)

(E) No baseline liver injury was observed by serum levels of ALT and AST.

(F) LW/BW (+/-SEM) was lower in EC-Wnt2-KO and EC-Wnt9b-KO mice, and even lower in EC-Wnt2-9b-DKO mice. (ns = not significant, *P < 0.05, **P < 0.01)

**Fig. S10 Workflow of single-cell transcriptomics analysis of Molecular Cartography data of EC-Wnt2-9b-DKO mice.** (Related to Figure 3 and STAR Methods)

See Methods for details.

**Fig. S11 Line plots showing changes of zonated genes between control and EC-Wnt2-9b-DKO mice.** (Related to Figure 3)

(A) Pericentral genes that decreased in EC-Wnt2-9b-DKO mice.

(B) Pericentral genes that increased in EC-Wnt2-9b-DKO mice.

(C) Periportal genes that had decreased zone 1 gene expression in EC-Wnt2-9b-DKO mice.

(D) Periportal genes that had similar or unchanged zone 1 gene expression in EC-Wnt2-9b-DKO mice.

(E) Periportal genes that had increased zone 1 gene expression in EC-Wnt2-9b-DKO mice. The hue around line plots represents SEM.

**Fig. S12 Workflow of single-cell transcriptomics analysis of Molecular Cartography data of FL6.13-treated mice.** (Related to Figure 6 and STAR Methods)

See Methods for details.

**Fig. S13 Feature plots showing expression of genes in control and FL6.13-treated animal.** (Related to Figure 6)

Blue: Control IgG; Red: FL6.13 treatment

**Fig. S14 Expression level of pericentral zonated genes in control and FL6.13-treated liver.** (Related to Figure 6)

(A) Violin plots showing expanded expression of Wnt target genes after FL6.13 treatment.

(B) Violin plots showing genes that were not increased after FL6.13 treatment.

**Fig. S15 Expression level of periportal zonated genes in control and FL6.13-treated liver.** (Related to Figure 6)

(A) Violin plots showing that expression of periportal genes was not decreased after FL6.13 treatment.

(B) Study design of control IgG and FL6.13 treatment for regeneration study.

**Fig. S16 Early treatment with FL6.13 worsens liver injury after acetaminophen overdose.** (Related to Figure 7)

(A) Study design showing dosing schedule of pan-FZD agonist FL6.13 or isotype control IgG administration. Single dose of 5 mg/kg control IgG or FL6.13 was i.p. administrated to mice at 12 hours post 600 mg/kg i.p. APAP injection. Mice were sacrificed at 48 hours for analysis.

(B) Gross images of livers showing necrosis and congestion in control IgG and FL6.13-treated animals at 48 hours.

(C) Serum levels of ALT, AST, direct bilirubin, total bilirubin, and ALP at 48 hours showing increased liver injury when FL6.13 was given at 12 hours. (The bars represent means ± SEM. ns = not significant, *P < 0.05)

(D) Representative IHC showing necrotic areas by HE, cell proliferation by Cyclin D1 and Ki67, and immune cell infiltration by CD45.

## Supplementary tables

**Table S1 Sequence of genotyping primers** (related to STAR Methods)

**Table S2 Size of PCR products** (related to STAR Methods)

**Table S3 Probe list for Molecular Cartography™** (related to STAR Methods)

**Table S4 Sequence of qPCR primers** (related to STAR Methods)

## Methods

### Animals

All animal husbandry and experimental procedures, including animal housing and diet, were performed under the guidelines and approval of the National Institutes of Health and the Institutional Animal Care and Use Committee at the University of Pittsburgh. Mice were fed regular chow in standard caging and kept under a 12-hour light-dark cycle with no enrichment. Lyve1-cre mice were purchased from Jackson Laboratories. *Wnt9b*^flox/flox^ mice were reported before (Carroll et al., 2005). *Wnt2* exon-2-floxed mice were generated by insertion of LoxP sites flanking exon 2, through use of CRISPR/Cas9 technology as described elsewhere (Pelletier et al., 2015). Briefly, using single-guide RNAs (sgRNAs), double strand breaks flanking exon 2 were targeted to optimal CAS9 target sites within introns 1 and 2. A synthetic floxed allele was generated as a single-stranded-oligodeoxynucleotide (ssODN) template for homology-directed repair (HDR). The HDR template was a megamer, 1166bp ssDNA oligo (Integrated DNA Technologies), encoding exon 2 flanked by LoxP sites with adjacent EcoR1 sites for genotyping and 5’ and 3’ homology arms homologous to the genomic sequences up-and down-stream of the 5’ and 3’ CAS9 cut sites in Introns 1 and 2. Homologous recombination was achieved by microinjection of a mixture of Cas9 protein (0.3 μM), I1 & I2 sgRNAs (21.23 ng/μl each) and the ssODN (10 ng/μl) into the pronuclei of fertilized embryos (C57BL/6J, The Jackson Laboratory). The injected zygotes were cultured overnight, the next day the embryos that developed to the 2-cell stage were transferred to the oviducts of pseudopregnant CD1 female recipients. Offspring were genotyped by PCR and RFLP analysis to confirm insertion of 5’and 3’ loxP sites. Founder mice, M14 and M15, were backcrossed with C57BL/6J wild-type (WT) mice for two generations and selected for the presence of *Wnt2*^flox/flox^ allele to generate N2 mice, which were then bred with *Wnt9b*^flox/flox^, Rosa-stop^flox/flox^-EYFP, and Lyve1-cre^+/-^ mice. Genotyping primers and PCR products are listed in Table S1 and S2.

Lyve1-cre^+/-^; *Wnt2*^flox/-^ *Wnt9b*^flox/flox^ Rosa-stop^flox/flox^-EYFP mice are hereafter referred to as EC-Wnt9b-KO. Lyve1-cre^+/-^; *Wnt2*^flox/flox^ *Wnt9b*^flox/-^ Rosa-stop^flox/flox^-EYFP mice are hereafter referred to as EC-Wnt2-KO. Lyve1-cre^+/-^; *Wnt2*^flox/flox^ *Wnt9b*^flox/flox^ Rosa-stop^flox/flox^-EYFP mice are hereafter referred to as EC-Wnt2-9b-DKO. The rest of mice were used as littermate controls (Control). EC-Wls-KO mice were described before (Preziosi et al., 2018). LRP5-6-LDKO mice were generated by i.p. injection of 1 x 10^12^ genome copies of AAV8-TBG-Cre (addgene, 107787-AAV8) per mice to LRP5-6 double floxed mice described before (Yang et al., 2014).

### Two-third or partial hepatectomy (PH)

Two to three-month-old male and female Control, EC-Wnt2-KO, EC-Wnt9b-KO, EC-Wnt2-9b-DKO mice were subjected to PH as describe before(Tan et al., 2006). Animals were sacrificed at 40 hours post-PH. Serum and liver tissues were harvested for further analysis.

### FL6.13 treatment

Two to three-month-old male wildtype, EC-Wnt2-9b-DKO, EC-Wls-KO, LRP5-6-LDKO mice were treated with 5 mg/kg control IgG or FL6.13 every other day for one week followed by PH. Animals were sacrificed at 24 hours post-PH. 1 mg/ml BrdU was given in drinking water. Serum and liver tissues were harvested for further analysis.

### Acetaminophen study

8-week-old male C57/BL6 mice were ordered from Jackson lab. Mice were given food and water from 6 pm to 9 pm in dark room and then fasted from 9 pm to 9 am next day. 600 mg/kg APAP was intraperitoneal (i.p.) injected, and food were given back. 12 hours or 32 hours post APAP injection, mice were randomly grouped and were i.p. injected with 5 mg/kg control IgG or FL6.13 in 0.9% saline. Mice were sacrificed at 48 hours or 60 hours. Serum and liver tissues were collected for further analysis.

### scRNA sequencing data analysis

scRNA sequencing data were published before and were analyzed at: https://www.livercellatlas.org/index.php (Guilliams et al., 2022).

### Molecular Cartography™

Detailed protocol of tissue processing, probe design, imaging, signal segmentation and barcoding was discussed before (Lotto et al., 2020). Probes used in this study are listed in Table S3. One animal per group was used. Molecular Cartography images were performed in ImageJ using genexyz Polylux tool plugin from Resolve BioSciences to examine specific Molecular Cartography signals.

Single-cell spatial transcriptomic analysis was performed according to the workflow in the Fig. S10 and Fig. S12. Two bioinformatic pipelines were applied to study the data. For the first pipeline, gene counts were quantified per cell based on the cell identification from QuPath software. Slide control, slide EC-Wnt2-9b-DKO, and slide FL6.13 treatment were integrated by R package *Seurat* (Hao et al., 2021). For quality control, cells with less than 10 gene-count were filtered out. Non-hepatocyte cells were defined by the cells with any expression of the five non-hepatocyte makers (*Lrat, Pecam1, Ptprc, Lyz2* and *Adgre1*) and were removed from further analysis when comparing control and FL6.13 treatment (Fig. S12). After pre-processing, dimension reduction method principal component analysis (PCA) and uniform manifold approximation and projection (UMAP) (McInnes et al., 2018) were performed on hepatocytes based on 16 zonated markers (pericentral markers: *Cldn2, Lect2, Glul, Oat, Cyp1a2, Cyp2e1, Gstm1, Rgn*; midzonal markers: *Pon1*; periportal markers: *Gls2, G6pc, Fbp1, Hsd17b13, Vtn, Cyp2f2, Pigr*). Eventually, cells were grouped into distinct clusters and annotated by the zonated genes. Feature plots and violin plots were generated by *Seurat* to visualize gene expression at cell-level and cluster-level.

For the second pipeline, a central line was drawn from pericentral to periportal bile duct based on landmark genes (*Cyp2e1, Glul*, and *Sox9*). Then an upper-bound and a lower-bound line were drawn parallel to the central line with 500-pixel extension. Bounded by these two lines, perpendicular lines were drawn to separate the region between pericentral and periportal into 9 zones. In total, 7, 9, and 9 pericentral-to-periportal regions were identified in slide control, slide EC-Wnt2-9b-DKO, and slide FL6.13 treatment, respectively. Gene counts and expression density (gene counts per area) were quantified within each zone and averaged across the defined pericentral-to-periportal regions. Cells located in these 9-zone areas were also identified based on their positions in the slides and traced back to the UMAPs to show the histological location of cells.

### RNA isolation and qPCR

Whole liver was homogenized in TRIzol™ (Thermo Scientific, 15596026) and nucleic acid was isolated through phenol-chloroform extraction. RNA was reverse transcribed into cDNA using SuperScript^®^ III (Invitrogen, 18080-044). Real-time PCR was performed in technical duplicate on a StepOnePlus™ Real-Time PCR System (Applied Biosystems, 4376600) using the Power SYBR^®^ Green PCR Master Mix (Applied Biosystems, 4367660). Target gene expression was normalized to housekeeping genes *Rn18s*, and fold change was calculated utilizing the ΔΔ-Ct method. Primers are listed in Table S4.

### Protein isolation and western blot

Snap frozen liver samples were homogenized in RIPA buffer with fresh proteinase and phosphatase inhibitor. The concentration of the protein was determined by the bicinchoninic acid assay. Protein sample was prepared with loading buffer (Bio-Rad, 1610737) with 5% 2-Mercaptoethanol (Bio-Rad, 161-0710) and subjected to electrophoresis. Protein sample was separated on pre-cast 7.5% or 4-20% polyacrylamide gels (Bio-Rad) and transferred to the PVDF membrane using the Trans-Blot Turbo Transfer System (Bio-Rad). Membranes were stained with Ponceau-S and blocked for 30 minutes with 5% nonfat dry milk (Cell signaling, 9999) or 5% BSA in Blotto buffer (0.15M NaCl, 0.02M Tris pH 7.5, 0.1% Tween in dH2O), and incubated with primary antibodies at 4°C overnight at the following concentrations: GS (Sigma, G2781, 1:2000), CYP2E1 (Sigma, HPA009128, 1:1000), CYP1A2 (Santa Cruz Biotechnology, sc-53241, 1:1000), Cyclin D1 (abcam, ab134175, 1:1000). Membranes were washed in Blotto buffer and incubated with the appropriate HRP-conjugated secondary antibody for 60 minutes at room temperature. Membranes were washed with Blotto buffer, and bands were developed utilizing SuperSignal® West Pico Chemiluminescent Substrate (Thermo Scientific, 34080) and visualized by time-gradient autoradiography.

### Immunohistochemistry

Livers were fixed in 10% buffered formalin for 48-72 hours prior to paraffin embedding. Blocks were cut into 4μm sections, deparaffinized, and washed with PBS. For antigen retrieval, samples were microwaved for 12 minutes in pH=6 sodium citrate buffer (CD45), or in pH=9 Tris-EDTA buffer (BrdU), or were pressure cooked for 20 minutes in pH=6 sodium citrate buffer (CYP2E1, CYP1A2, Ki67, CYP2F2), or in pH=9 Tris-EDTA buffer (GFP, Cyclin D1, RGN, PIGR). For BrdU, slides were then incubated with 2M HCl for one hour at room temperature and washed with 0.5M Borax for 5 minutes. GS staining doesn’t need antigen retrieval. Samples were then placed in 3% H_2_O_2_ for 10 minutes to quench endogenous peroxide activity. After washing with PBS, slides were blocked for 10 minutes. The primary antibodies were incubated at the following concentrations in PBS: CD45 (Santa Cruz Biotechnology, sc-53665, 1:100), BrdU (Accurate Chemicals, OBT0030A, 1:75), GS (Sigma, G2781, 1:3000), CYP2E1 (Sigma, HPA009128, 1:100), CYP1A2 (Santa Cruz Biotechnology, sc-53241, 1:100), Ki67 (Cell signaling, 12202, 1:500), CYP2F2 (Santa Cruz Biotechnology, sc-374540, 1:100), GFP (Cell signaling, 2956, 1:100), Cyclin D1 (abcam, ab134175, 1:200), RGN (Santa Cruz Biotechnology, sc-390098, 1:100), PIGR (R&D Systems, AF2800, 1:100) for one hour at room temperature. Samples were washed with PBS three times and incubated with the appropriate biotinylated secondary antibody (Vector Laboratories) diluted 1:250 in antibody diluent for 15 minutes at room temperature. Samples were washed with PBS three times and sensitized with the Vectastain® ABC kit (Vector Laboratories, PK-6101). Following three washes with PBS color was developed with DAB Peroxidase Substrate Kit (Vector Laboratories, SK-4100), followed by quenching in distilled water. Slides were counterstained with hematoxylin (Thermo Scientific, 7211), dehydrated to xylene and coverslips applied with Cytoseal™ XYL (Thermo Scientific, 8312-4). Images were taken on a Zeiss Axioskop 40 inverted brightfield microscope.

### Immunofluorescence

For triple staining of CK19, CD31, and GFP, paraffin sections were deparaffinized and pressure cooked in pH=9 Tris-EDTA buffer for 20min. Slides were permeabilized with 0.3% Triton X-100 in PBS for 20 minutes at room temperature and then blocked with 5% normal donkey serum in 0.3% Triton X-100 in PBS (antibody diluent) for 30 minutes at room temperature. Antibodies were diluted as follows: CK19 (DSHB, TROMA-III, 1:10), CD31 (BD, AF3628, 1:100), GFP (Cell signaling, 2956, 1:100), in antibody diluent and incubated at 4°C overnight. Samples were washed three times in 0.1% Triton X-100 in PBS (wash solution) and incubated with the proper fluorescent secondary antibody (AlexaFluor 488/555/647, Invitrogen) diluted 1:300 in antibody diluent for two hours at room temperature. Samples were washed three times and incubated with DAPI (Sigma, B2883) for 1 minute. Samples were washed three times and mounted with gelvator. Images were taken on a Nikon Eclipse Ti epifluorescence microscope or a Zeiss LSM700 confocal microscope and were analyzed with Image J.

### Statistics

Statistical comparison between two groups was done with the unpaired Student’s t test. Multiple-group comparison was done with one-way analysis of variance (ANOVA). GraphPad Prism version 9.0 was used for graph generation, and a P value of less than 0.05 was considered significant. The bars represent means ± SEM. ns = not significant, *P < 0.05, **P < 0.01, ***P < 0.001, and ****P < 0.0001.

## Notes

**Funding:** This work was supported by NIH grants 1R01CA251155, 1R01CA204586, 1R01DK62277, 1R01DK116993, and Endowed Chair for Experimental Pathology to SPM, and by NIH grant 1P30DK120531 to Pittsburgh Liver Research Center (PLRC) for services provided by the Genomics and Systems Biology Core. This research was also supported in part by the University of Pittsburgh Center for Research Computing through the resources provided. This work was also supported by the University of Toronto Medicine by Design program (grant# 72058117), which receives funding from the Canada First Research Excellence Fund (to S.S and S.A).

