## Supplementary figures and images for "Dynamic control of metabolic zonation and liver repair by endothelial cell Wnt2 and Wnt9b revealed by single cell spatial transcriptomics using Molecular Cartography"

### Figure S1.tif

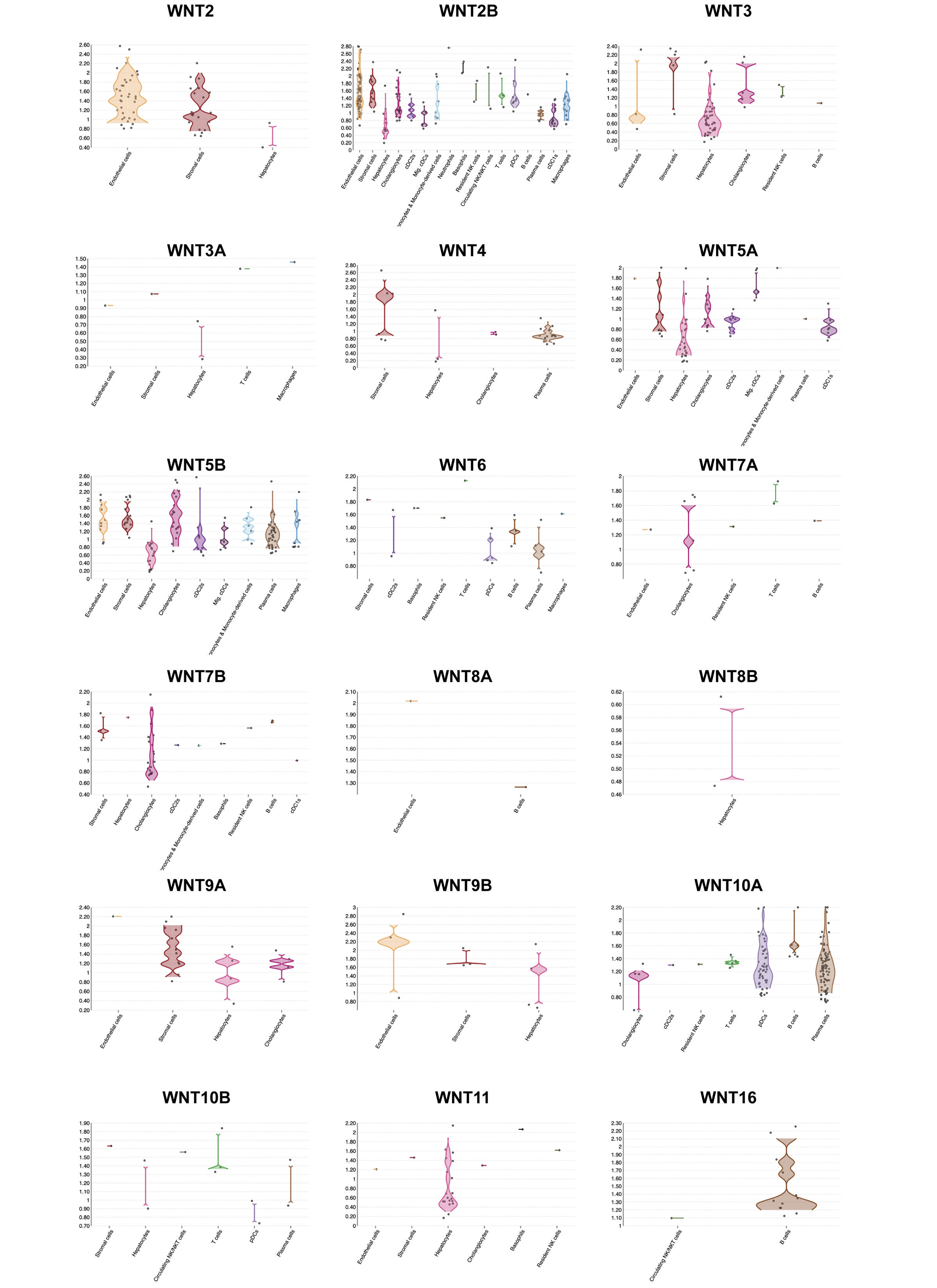

### Figure S2.tif

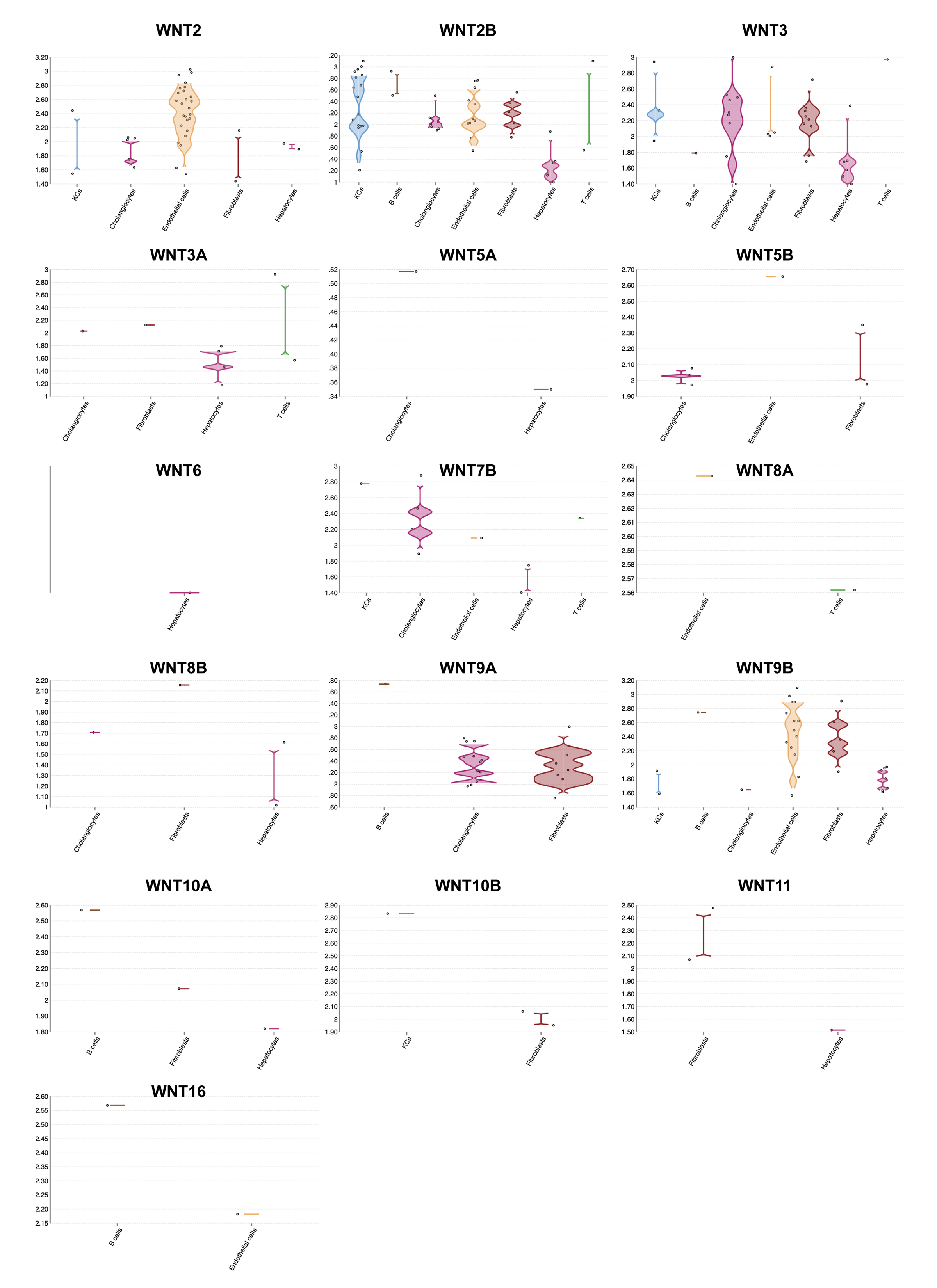

### Figure S3.tif

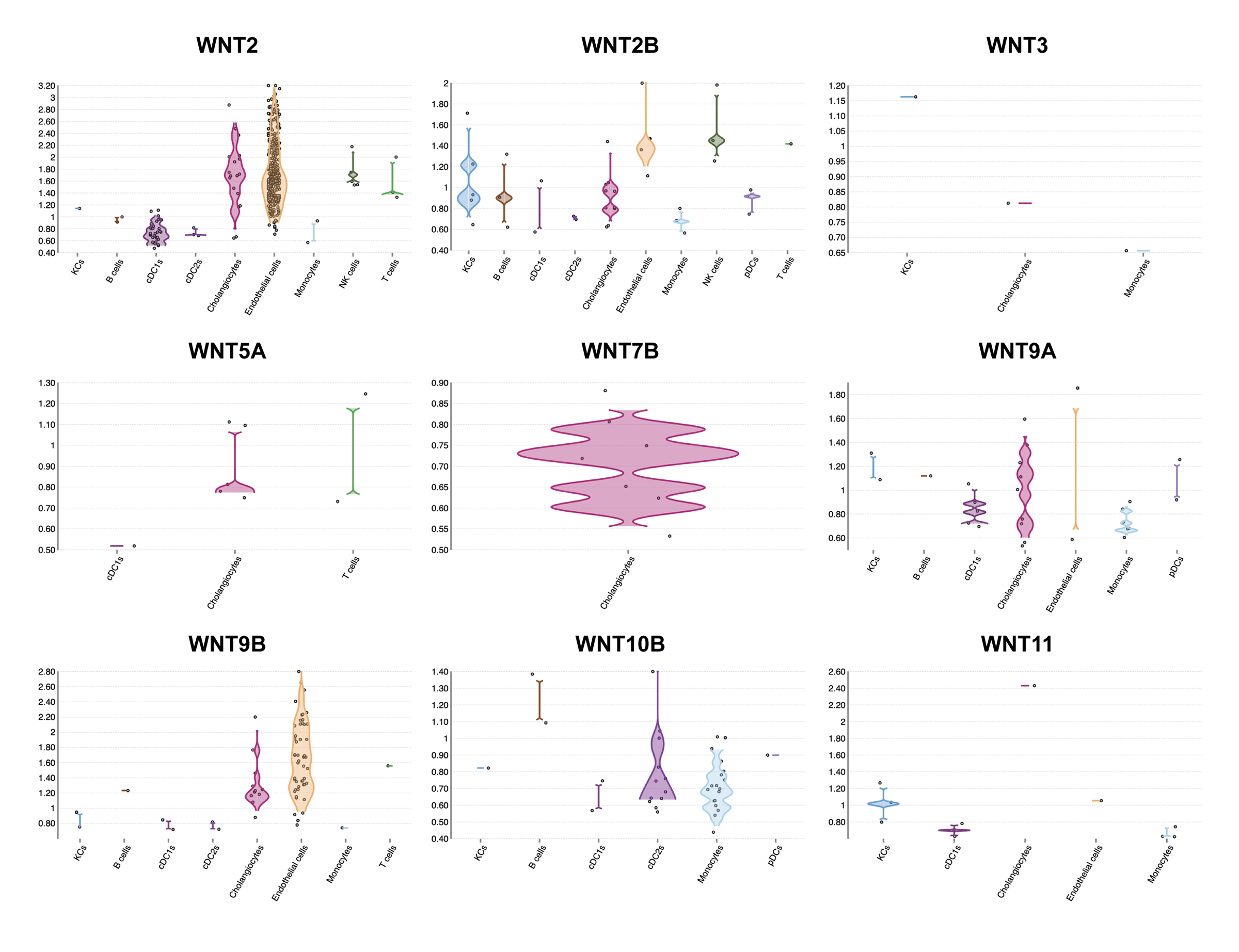

### Figure S4.tif

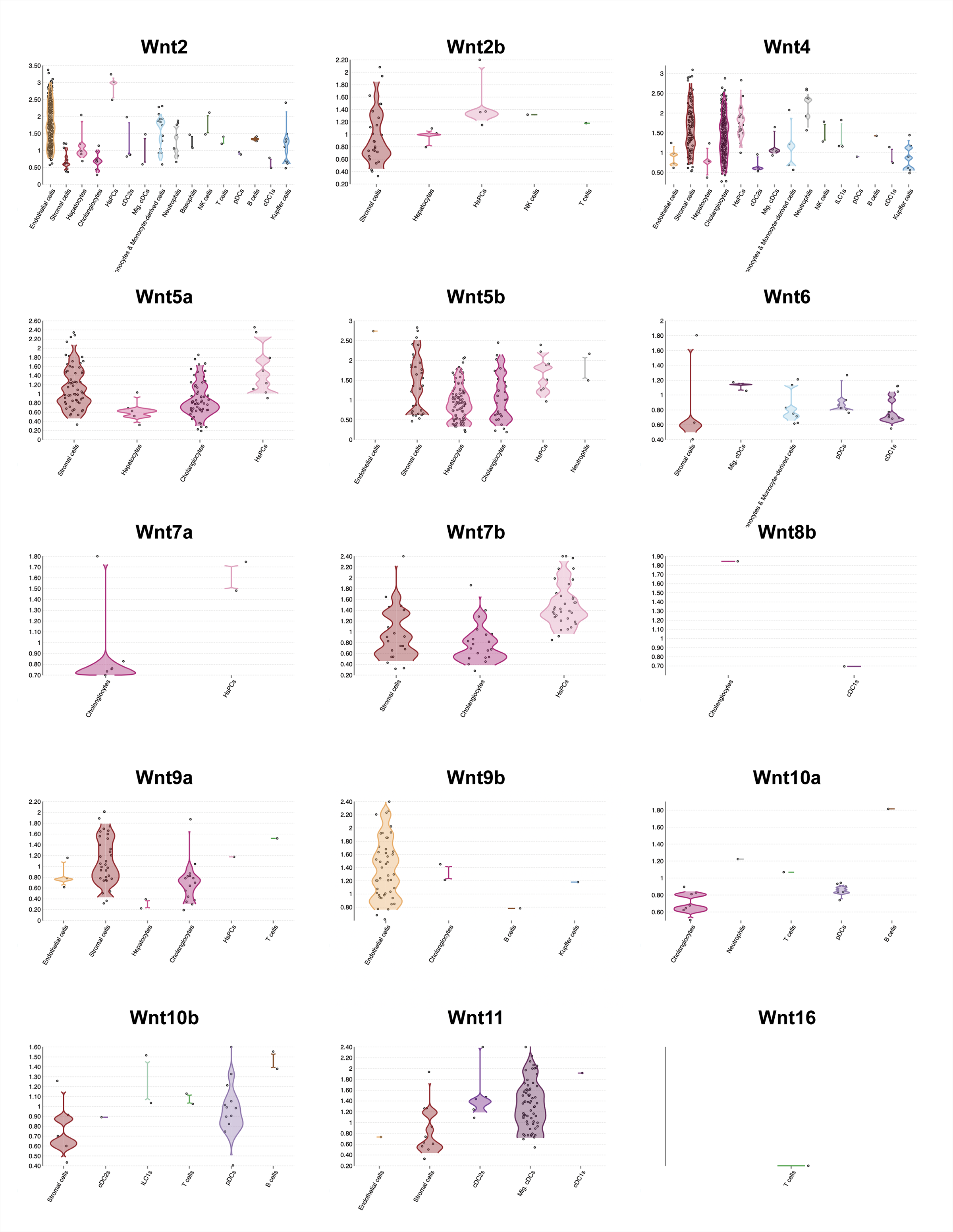

### Figure S5.tif

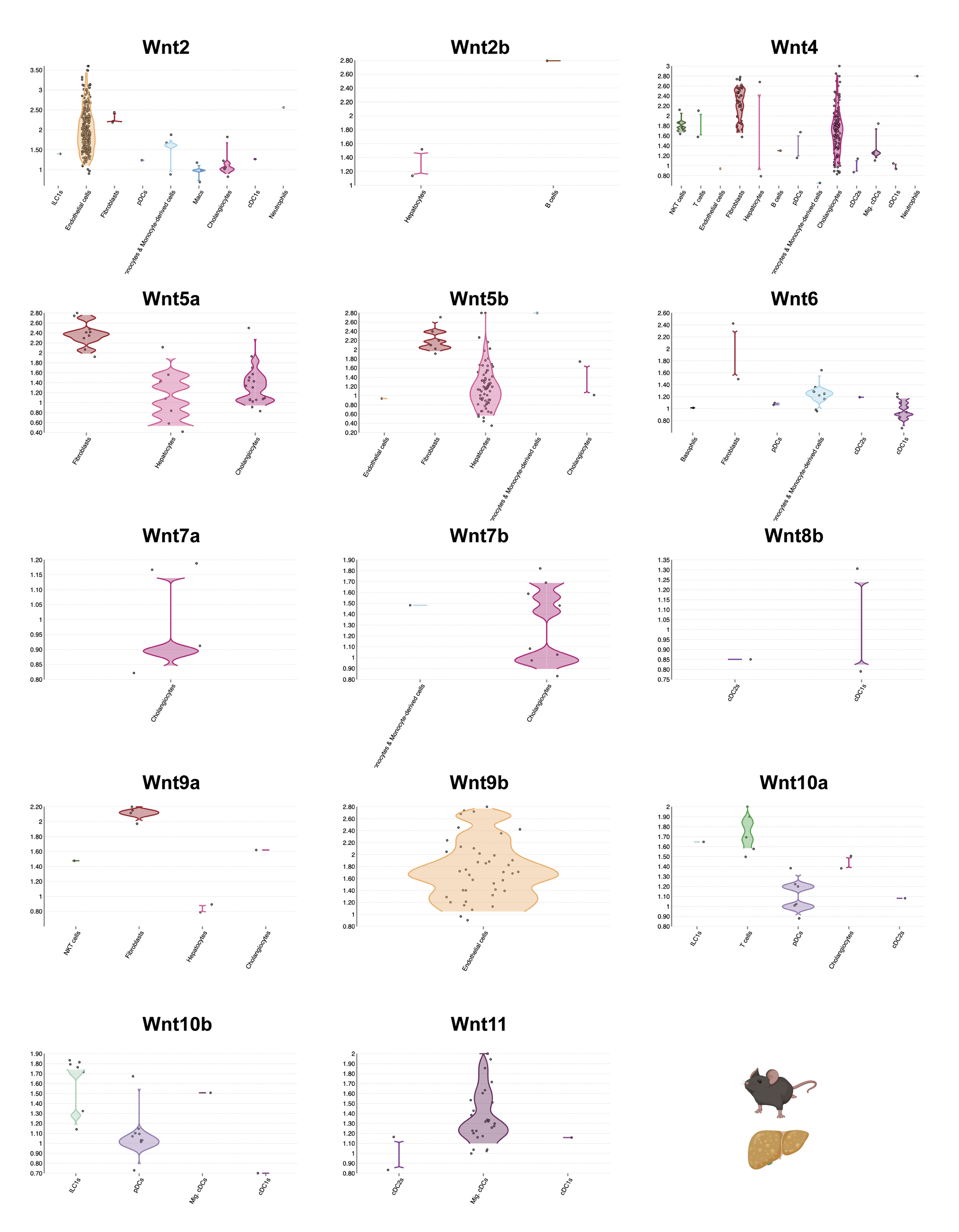

### Figure S6.tif

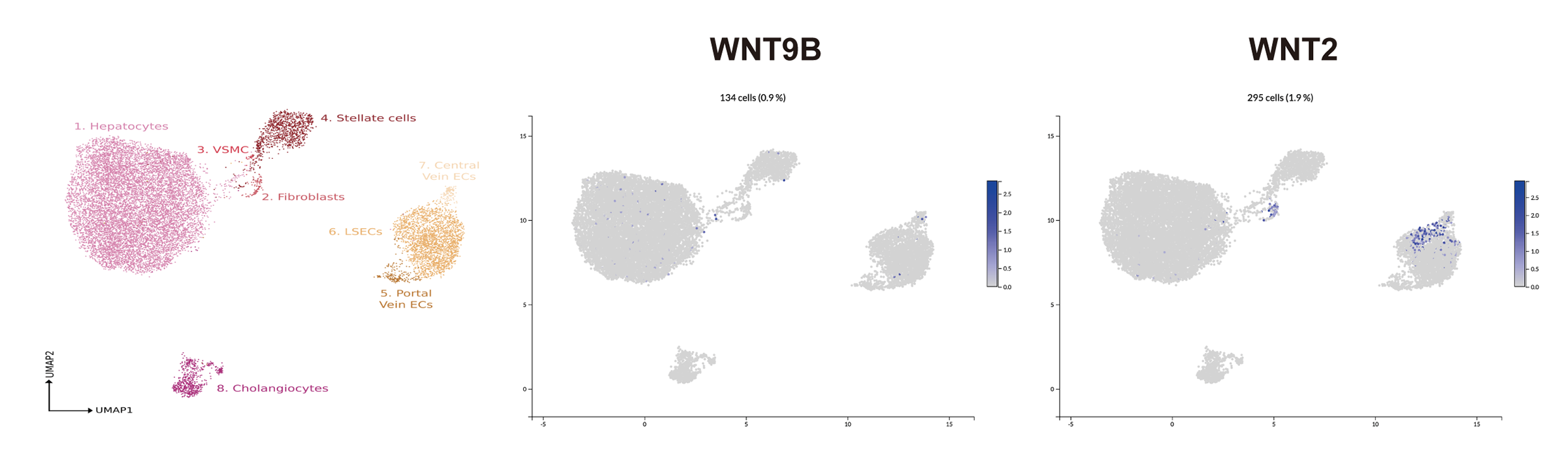

### Figure S7.tif

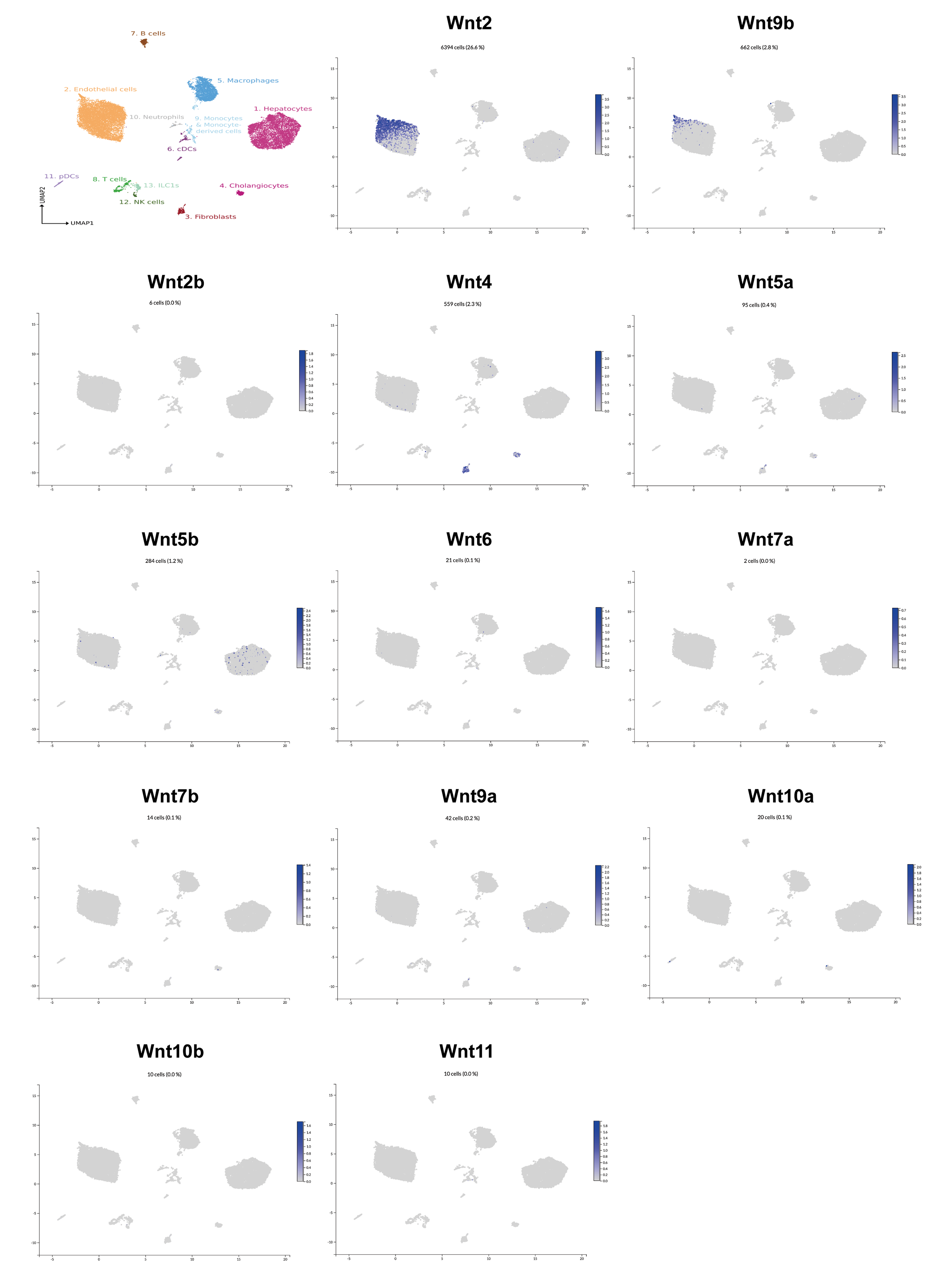

### Figure S8.tif

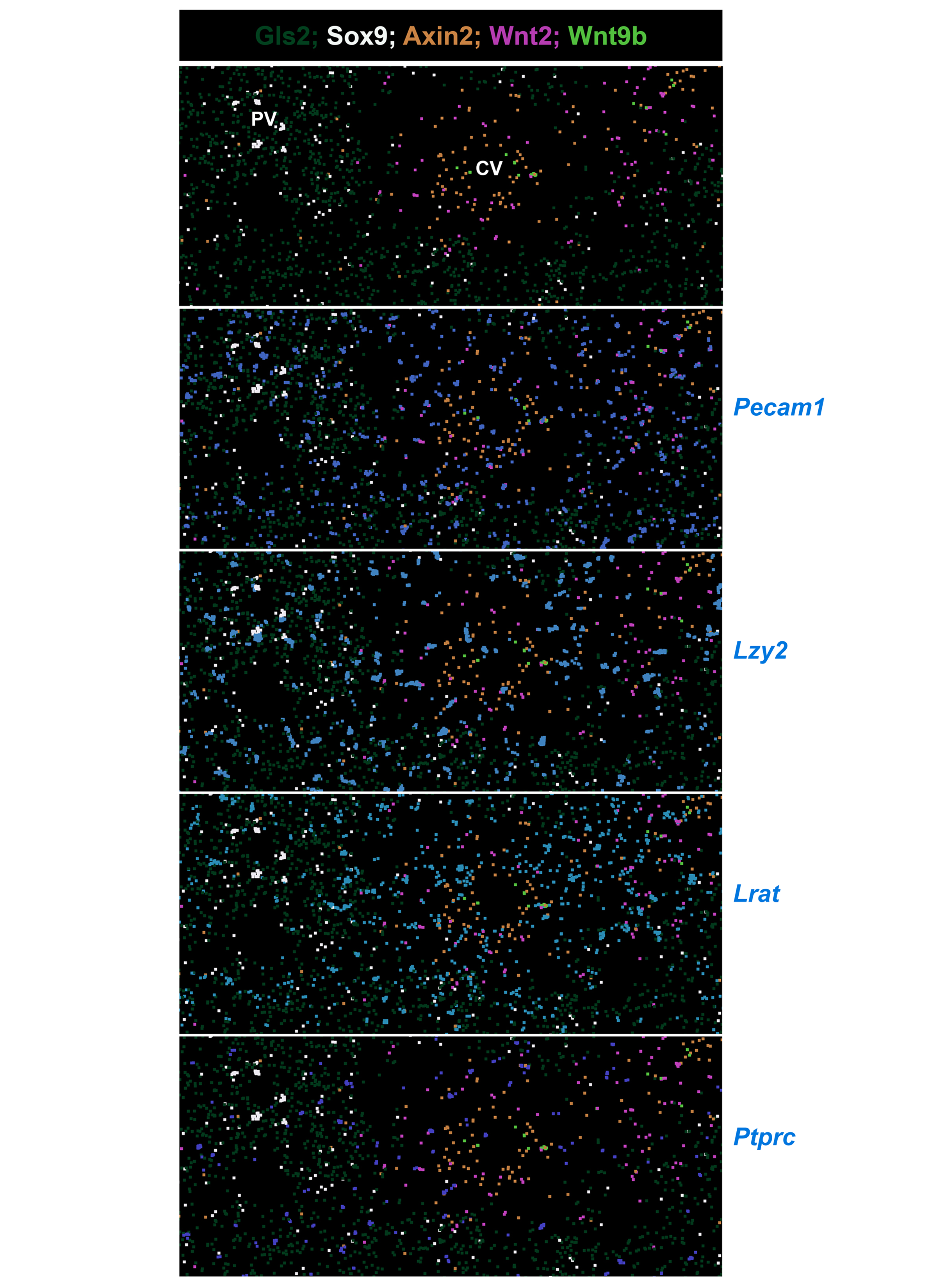

### Figure S9.tif

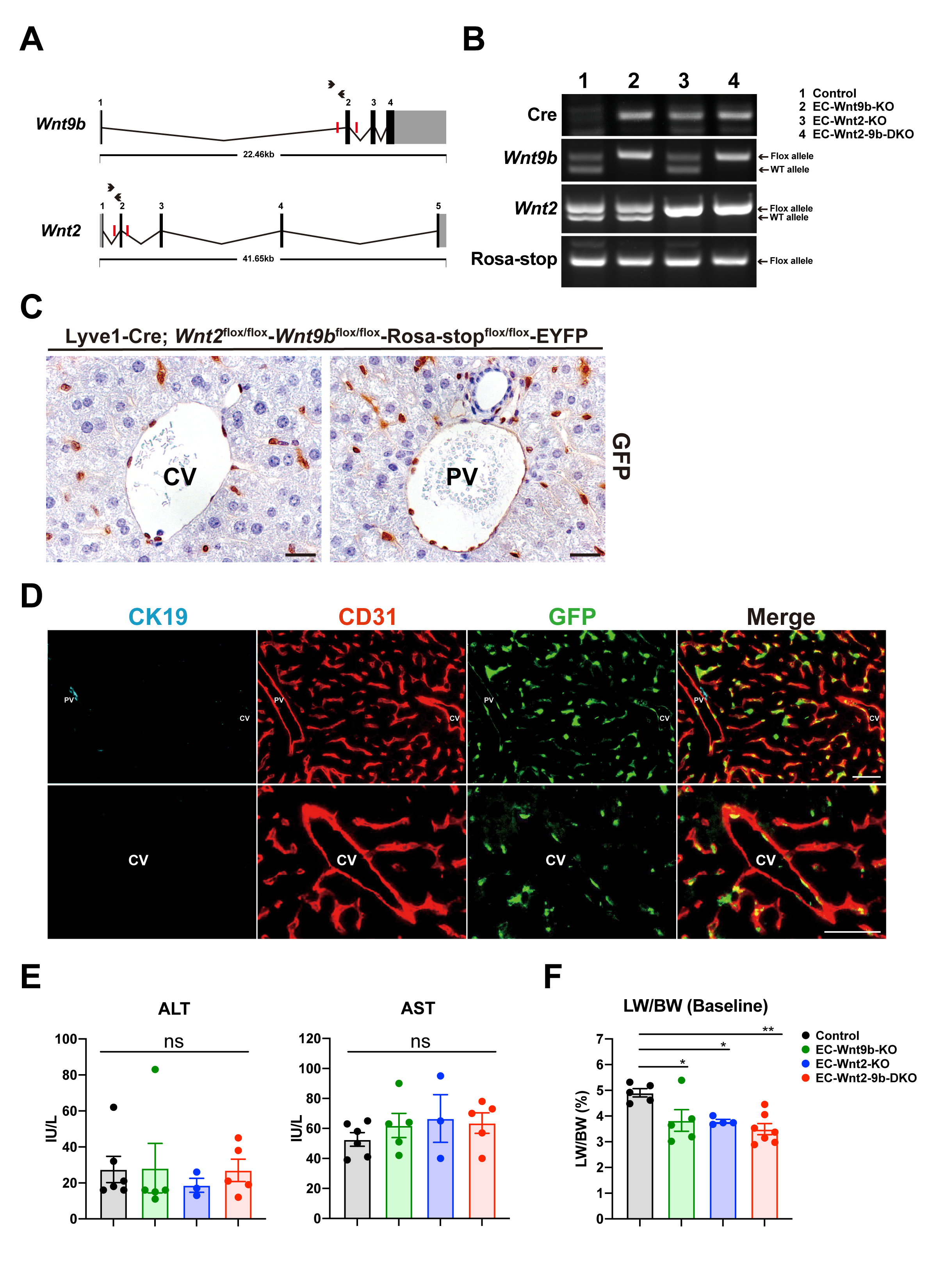

### Figure S10.tif

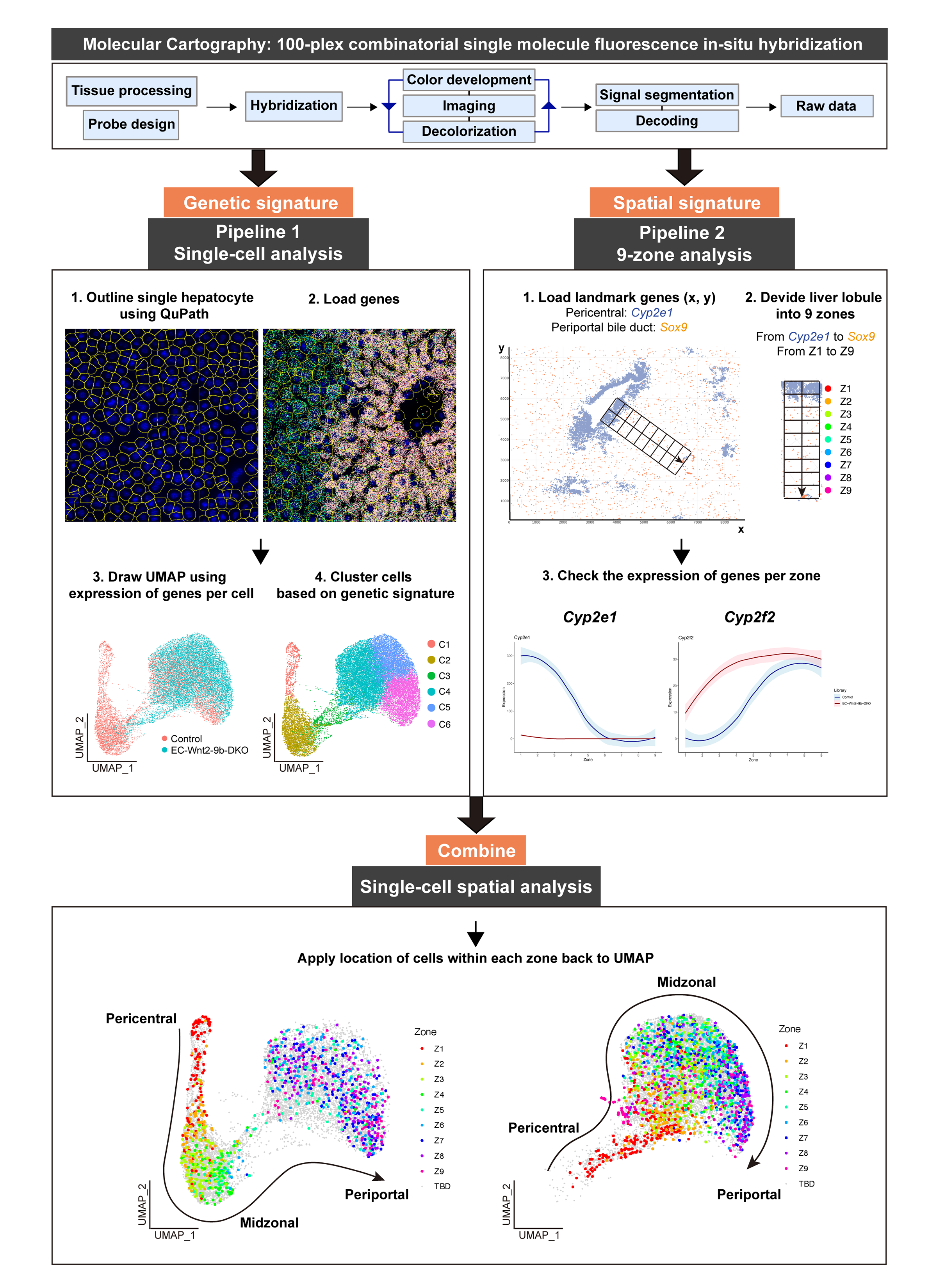

### Figure S11.tif

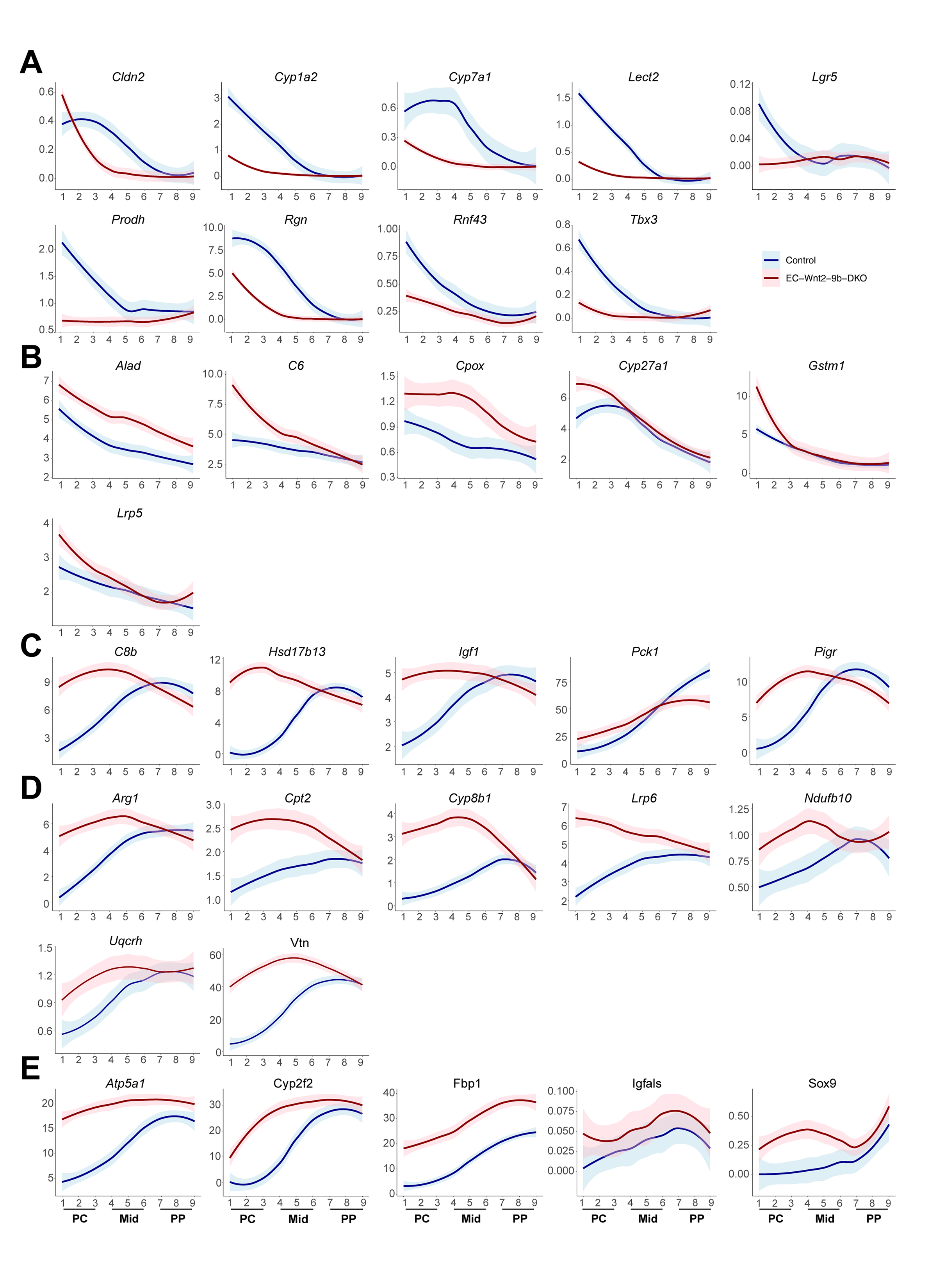

### Figure S12.tif

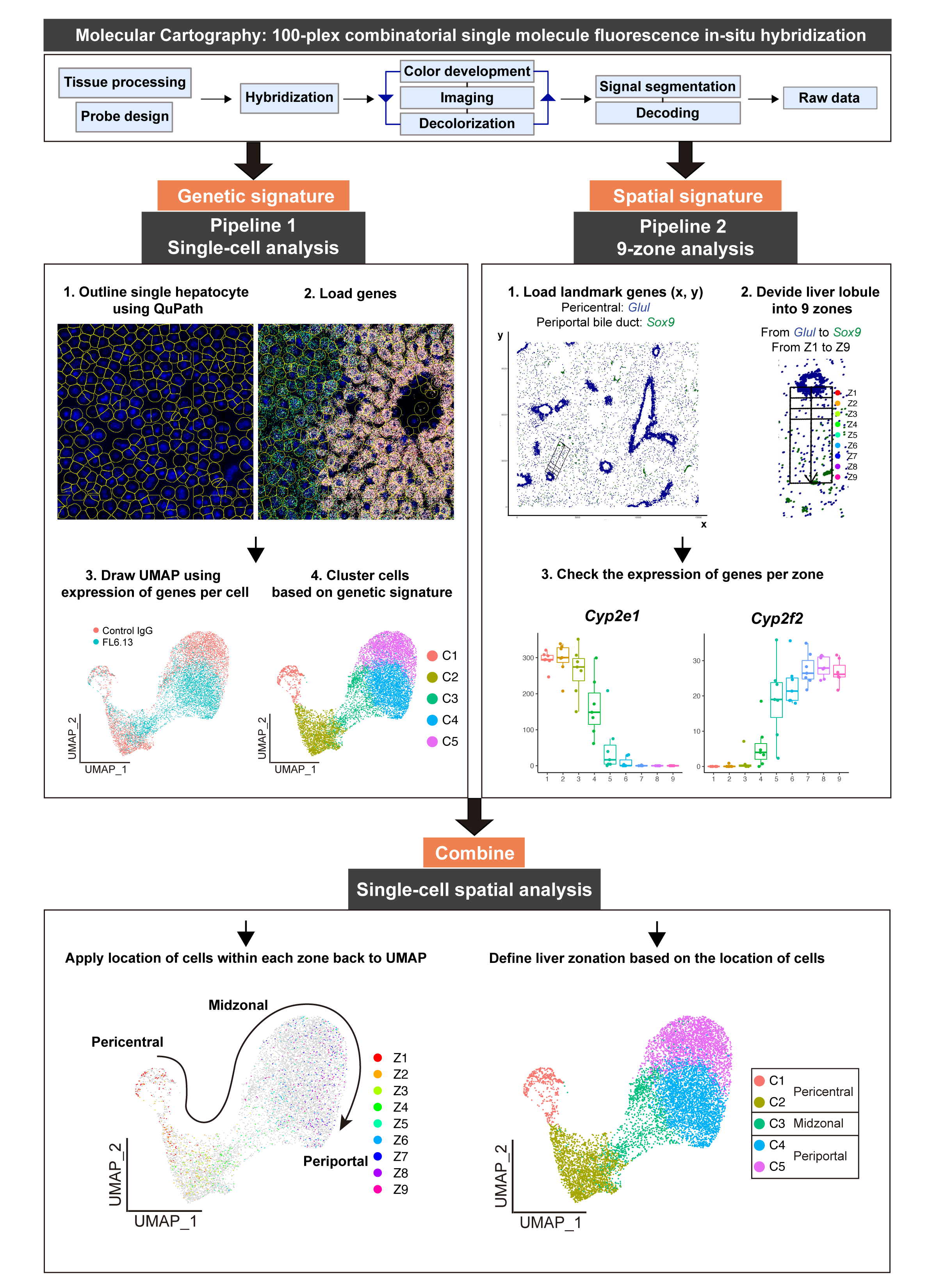

### Figure S13.tif

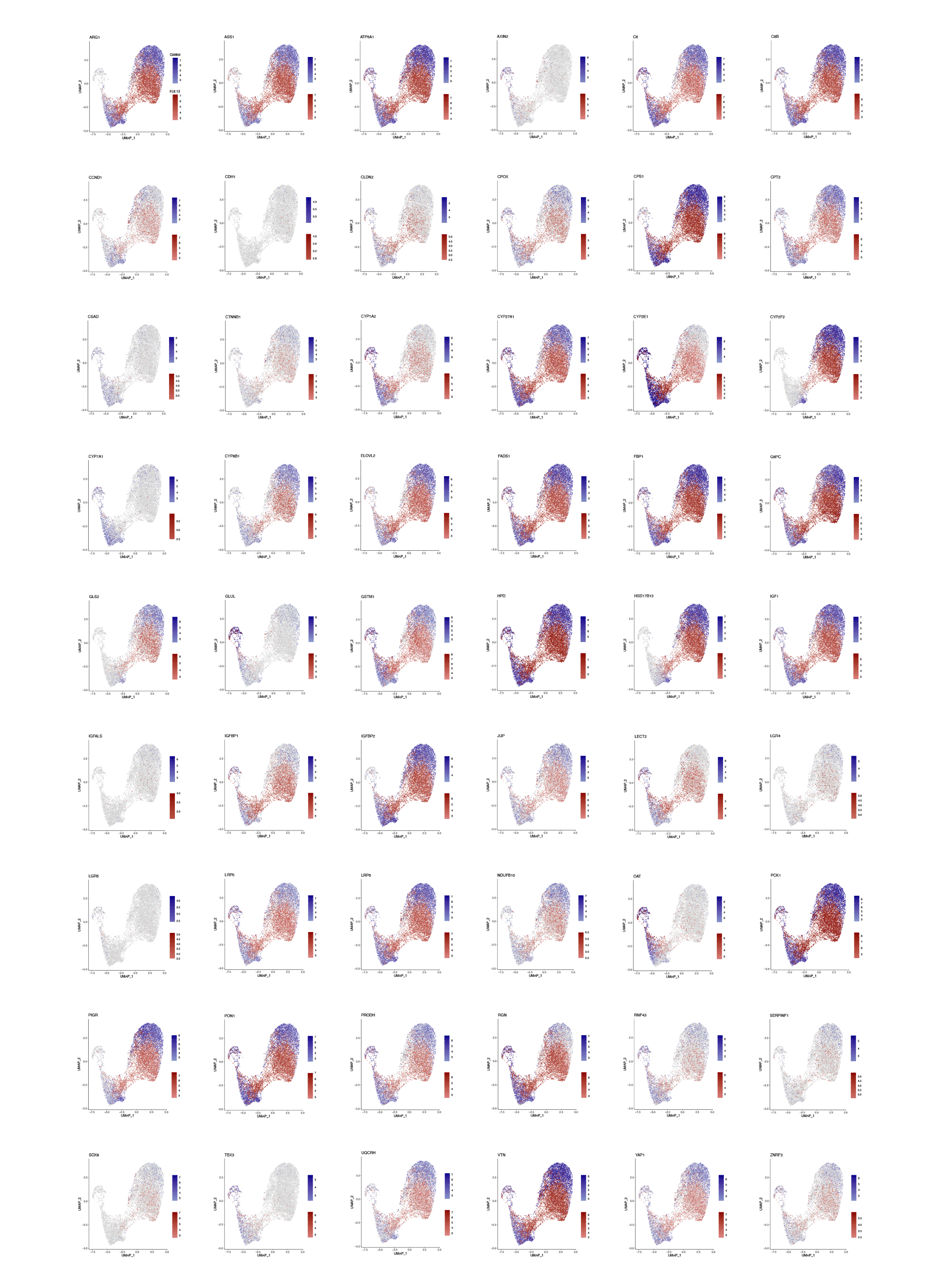

### Figure S14.tif

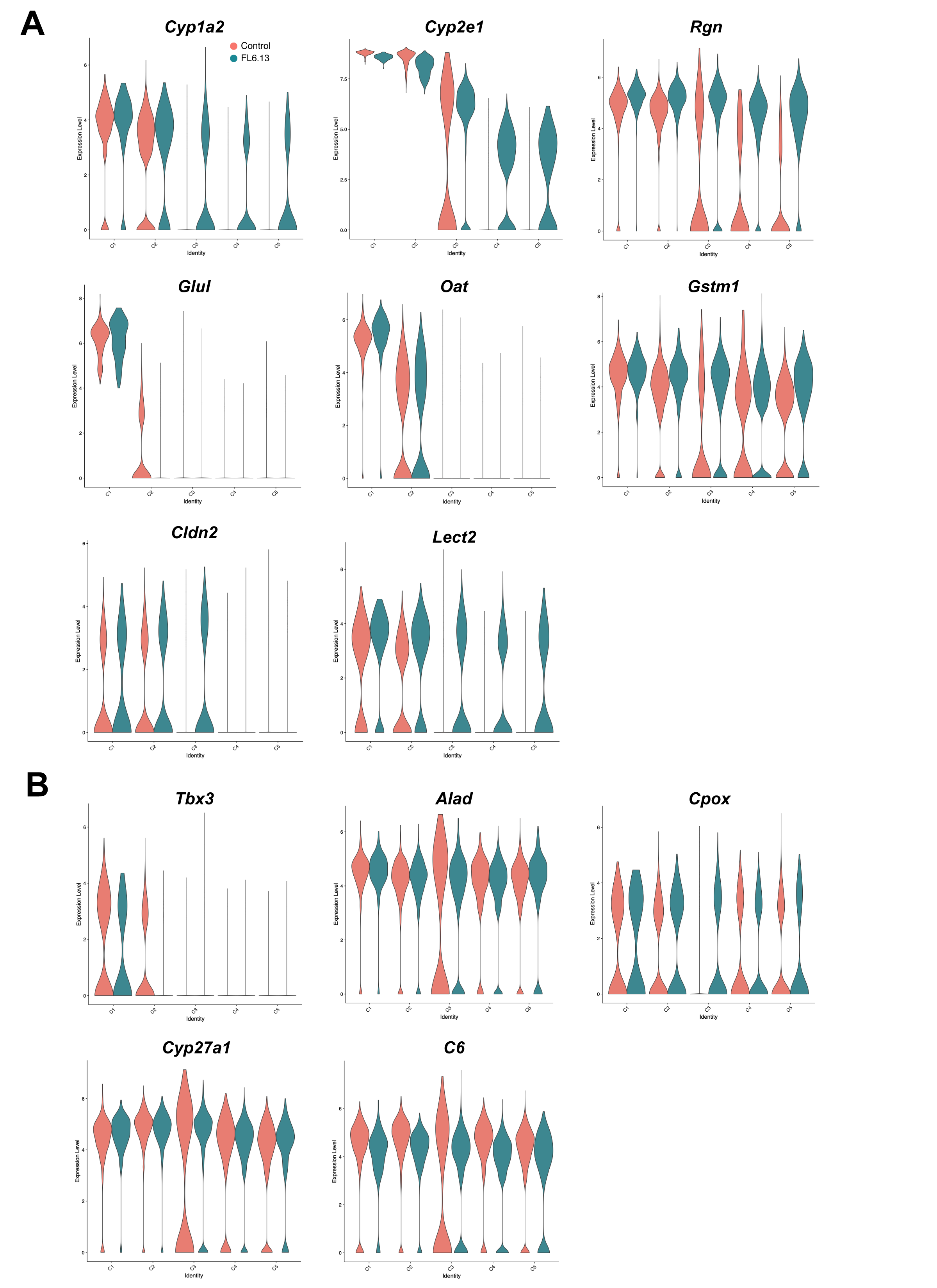

### Figure S15.tif

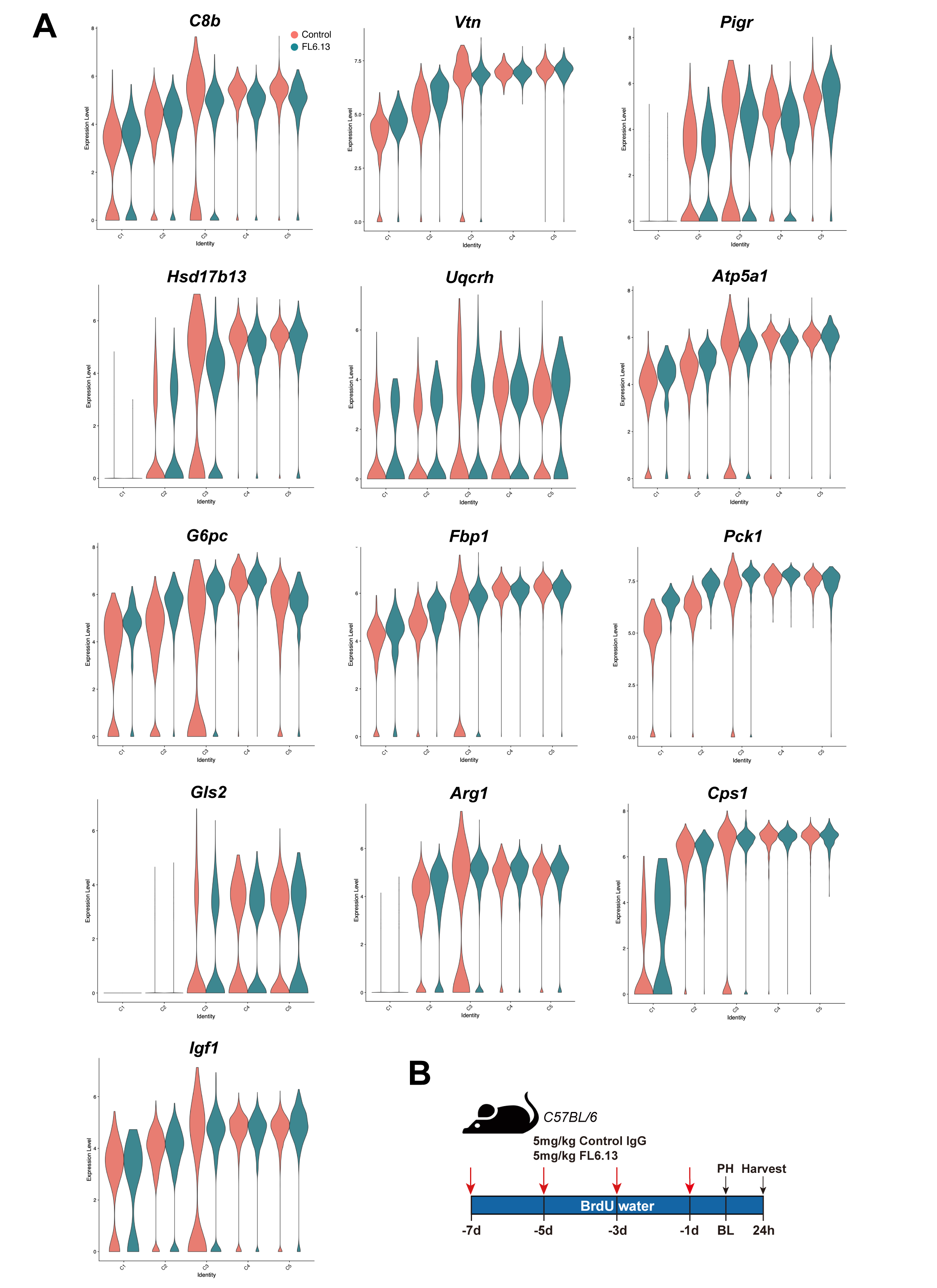

### Figure S16.tif

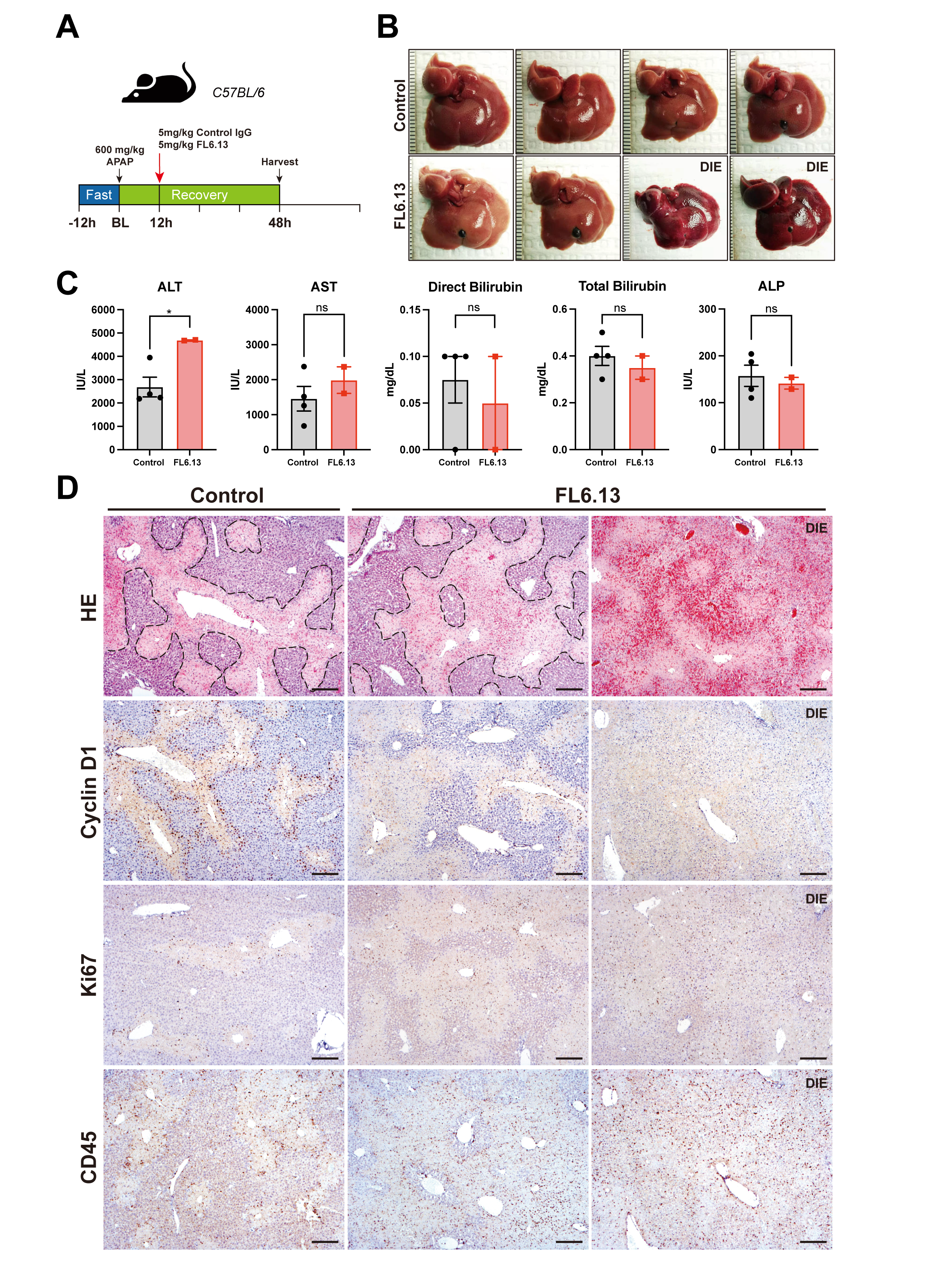
