## Supplementary table for "Dynamic control of metabolic zonation and liver repair by endothelial cell Wnt2 and Wnt9b revealed by single cell spatial transcriptomics using Molecular Cartography": Table S1 .docx

**Table S1 Sequence of genotyping primers** (related to STAR Methods)

| Primer name | Sequence (5’-3’) |
| --- | --- |
| Lyve1-Cre-F_wt | TGCCACCTGAAGTCTCTCCT |
| Lyve1-Cre-F_mutant | GAGGATGGGGACTGAAACTG |
| Lyve1-Cre-R | TGAGCCACAGAAGGGTTAGG |
| ROSA-EYFP-F_wt | GGAGCGGGAGAAATGGATATG |
| ROSA-EYFP-F_mutant | AAGACCGCGAAGAGTTTGTC |
| ROSA-EYFP-R | AAAGTCGCTCTGAGTTGTTAT |
| Wnt9b-F | GCAGAATCTGGAGAACTTGGC |
| Wnt9b-R | GTGAGAAGGAAGATGGTGAGC |
| Wnt2-F | CCCAGCAGGTGCTAAGAGG |
| Wnt2-R | CAATGGCACGCATCACATCT |
