## Supplementary table for "Dynamic control of metabolic zonation and liver repair by endothelial cell Wnt2 and Wnt9b revealed by single cell spatial transcriptomics using Molecular Cartography": Table S2.docx

**Table S2 Size of PCR products** (related to STAR Methods)

|  | WT (bp) | Mutant (bp) |
| --- | --- | --- |
| Lyve1-Cre | 425 | 210 |
| Rosa-EYFP | 600 | 324 |
| Wnt9b | 219 | 300 |
| Wnt2 | 540 | 592 |
