## Supplementary table for "Dynamic control of metabolic zonation and liver repair by endothelial cell Wnt2 and Wnt9b revealed by single cell spatial transcriptomics using Molecular Cartography": Table S3 .docx

**Table S3 Probe list for Molecular Cartography** (related to STAR Methods)

| **Item** | **CatNo** | **Species** | **Design Target ID** | **Gene Name** |
| --- | --- | --- | --- | --- |
| 1 | P0D1P | Mus musculus | ENSMUSG00000006932 | Ctnnb1 |
| 2 | P0DCL | Mus musculus | ENSMUSG00000015957 | Wnt11 |
| 3 | P0ECZ | Mus musculus | ENSMUSG00000002588 | Pon1 |
| 4 | P1DC0 | Mus musculus | ENSMUSG00000029671 | Wnt16 |
| 5 | P2D1R | Mus musculus | ENSMUSG00000024913 | Lrp5 |
| 6 | P2DC1 | Mus musculus | ENSMUST00000054294 | Fzd1 |
| 7 | P2Y49 | Mus musculus | ENSMUSG00000000303 | Cdh1 |
| 8 | P3DC2 | Mus musculus | ENSMUSG00000022297 | Fzd6 |
| 9 | P3EC1 | Mus musculus | ENSMUST00000102840 | Ass1 |
| 10 | P3F4T | Mus musculus | ENSMUSG00000004730 | Adgre1 |
| 11 | P4DC3 | Mus musculus | ENSMUST00000114246 | Fzd7 |
| 12 | P4EC2 | Mus musculus | ENSMUST00000020161 | Arg1 |
| 13 | P5DC4 | Mus musculus | ENSMUST00000041080 | Fzd8 |
| 14 | P5EC3 | Mus musculus | ENSMUST00000045602 | Ndufb10 |
| 15 | P638C | Mus musculus | ENSMUST00000092163 | Lyz2 |
| 16 | P6787 | Mus musculus | ENSMUST00000019469 | G6pc |
| 17 | P6C1X | Mus musculus | ENSMUSG00000020053 | lgf1 |
| 18 | P6DC5 | Mus musculus | ENSMUST00000032322 | Lrp6 |
| 19 | P7DC6 | Mus musculus | ENSMUSG00000050199 | Lgr4 |
| 20 | P7EC5 | Mus musculus | ENSMUST00000092888 | Fbp1 |
| 21 | P8DC7 | Mus musculus | ENSMUSG00000034177 | Rnf43 |
| 22 | P8EC6 | Mus musculus | ENSMUST00000050714 | lgfals |
| 23 | P948E | Mus musculus | ENSMUST00000003100 | Cyp2f2 |
| 24 | P9DC8 | Mus musculus | ENSMUSG00000041961 | Znrf3 |
| 25 | P9M7Y | Mus musculus | ENSMUST00000092623 | Rspo3 |
| 26 | PAN7Y | Mus musculus | ENSMUST00000044776 | Gls2 |
| 27 | PCN7Z | Mus musculus | ENSMUST00000029017 | Pck1 |
| 28 | PDECA | Mus musculus | ENSMUSG00000021364 | Elovl2 |
| 29 | PEECC | Mus musculus | ENSMUSG00000010663 | Fads1 |
| 30 | PEV7V | Mus musculus | ENSMUST00000029905 | Cyp7a1 |
| 31 | PFE89 | Mus musculus | ENSMUST00000117102 | Fzd10 |
| 32 | PFG43 | Mus musculus | ENSMUSG00000026395 | Ptprc |
| 33 | PFVCL | Mus musculus | ENSMUST00000057893 | Fzd2 |
| 34 | PGC15 | Mus musculus | ENSMUSG00000022382 | Wnt7b |
| 35 | PGECE | Mus musculus | ENSMUSG00000029656 | C8b |
| 36 | PGVC0 | Mus musculus | ENSMUSG00000001552 | Jup |
| 37 | PHE8C | Mus musculus | ENSMUST00000131309 | Fzd3 |
| 38 | PHVC1 | Mus musculus | ENSMUSG00000024182 | Axin1 |
| 39 | PJDCH | Mus musculus | ENSMUST00000024954 | Epas1 |
| 40 | PJE8D | Mus musculus | ENSMUST00000058755 | Fzd4 |
| 41 | PJM75 | Mus musculus | ENSMUST00000010941 | Wnt2 |
| 42 | PJR71 | Mus musculus | ENSMUST00000093962 | Ccnd1 |
| 43 | PJVC2 | Mus musculus | ENSMUSG00000005871 | Apc |
| 44 | PKCCK | Mus musculus | ENSMUST00000023734 | Wnt1 |
| 45 | PKDCJ | Mus musculus | ENSMUSG00000015522 | Arnt |
| 46 | PKE8E | Mus musculus | ENSMUSG00000045005 | Fzd5 |
| 47 | PKM76 | Mus musculus | ENSMUST00000018630 | Wnt9b |
| 48 | PKT6L | Mus musculus | ENSMUST00000000579 | Sox9 |
| 49 | PKVC3 | Mus musculus | ENSMUSG00000031169 | Porcn |
| 50 | PMA5F | Mus musculus | ENSMUSG00000028393 | Alad |
| 51 | PMCCM | Mus musculus | ENSMUST00000029429 | Wnt2b |
| 52 | PMDCK | Mus musculus | ENSMUST00000060077 | Cpox |
| 53 | PMECJ | Mus musculus | ENSMUST00000017488 | Vtn |
| 54 | PMP1L | Mus musculus | ENSMUSG00000039323 | lgfbp2 |
| 55 | PMVC4 | Mus musculus | ENSMUSG00000028173 | Wls |
| 56 | PNCCN | Mus musculus | ENSMUST00000000127 | Wnt3 |
| 57 | PNDCM | Mus musculus | ENSMUSG00000022181 | C6 |
| 58 | PNVC5 | Mus musculus | ENSMUSG00000018569 | Cldn7 |
| 59 | PPCCP | Mus musculus | ENSMUST00000010044 | Wnt3a |
| 60 | PPDCN | Mus musculus | ENSMUST00000054889 | Cldn2 |
| 61 | PPHCH | Mus musculus | ENSMUSG00000034528 | Hsd17b13 |
| 62 | PPVC6 | Mus musculus | ENSMUSG00000021539 | Lect2 |
| 63 | PQCCQ | Mus musculus | ENSMUST00000045747 | Wnt4 |
| 64 | PQE8J | Mus musculus | ENSMUST00000062572 | Fzd9 |
| 65 | PQVC7 | Mus musculus | ENSMUSG00000003526 | Prodh |
| 66 | PR47V | Mus musculus | ENSMUSG00000053110 | Yap1 |
| 67 | PR56S | Mus musculus | ENSMUSG00000018604 | Tbx3 |
| 68 | PR81H | Mus musculus | ENSMUSG00000020717 | Pecam1 |
| 69 | PRCCR | Mus musculus | ENSMUSG00000021994 | Wnt5a |
| 70 | PRVC8 | Mus musculus | ENSMUST00000034860 | Cyp1a2 |
| 71 | PS68V | Mus musculus | ENSMUST00000027144 | Cps1 |
| 72 | PSCCS | Mus musculus | ENSMUST00000006716 | Wnt6 |
| 73 | PSDCR | Mus musculus | ENSMUST00000031398 | Hpd |
| 74 | PSVC9 | Mus musculus | ENSMUST00000023832 | Rgn |
| 75 | PT86R | Mus musculus | ENSMUSG00000000753 | Serpinf1 |
| 76 | PTCCT | Mus musculus | ENSMUST00000032180 | Wnt7a |
| 77 | PTDCS | Mus musculus | ENSMUST00000084500 | Oat |
| 78 | PTF9N | Mus musculus | ENSMUST00000030687 | Rspo1 |
| 79 | PTVCA | Mus musculus | ENSMUST00000078676 | Uqcrh |
| 80 | PV85R | Mus musculus | ENSMUSG00000025428 | Atp5a1 |
| 81 | PVCCV | Mus musculus | ENSMUST00000012426 | Wnt8a |
| 82 | PVF4H | Mus musculus | ENSMUSG00000025479 | Cyp2e1 |
| 83 | PWCCW | Mus musculus | ENSMUST00000041163 | Wnt8b |
| 84 | PWDCV | Mus musculus | ENSMUSG00000019838 | Slc16a10 |
| 85 | PWF4J | Mus musculus | ENSMUSG00000020140 | Lgr5 |
| 86 | PWVCD | Mus musculus | ENSMUSG00000028607 | Cpt2 |
| 87 | PXCCX | Mus musculus | ENSMUSG00000000126 | Wnt9a |
| 88 | PXDCW | Mus musculus | ENSMUST00000062474 | Cyp8b1 |
| 89 | PXF4K | Mus musculus | ENSMUSG00000000142 | Axin2 |
| 90 | PXV8A | Mus musculus | ENSMUSG00000023044 | Csad |
| 91 | PXVCE | Mus musculus | ENSMUST00000027675 | Pigr |
| 92 | PXWCD | Mus musculus | ENSMUSG00000058135 | Gstm1 |
| 93 | PYCCY | Mus musculus | ENSMUST00000006718 | Wnt10a |
| 94 | PYDCX | Mus musculus | ENSMUST00000027356 | Cyp27a1 |
| 95 | PYWCE | Mus musculus | ENSMUST00000020704 | lgfbp1 |
| 96 | PZ78L | Mus musculus | ENSMUST00000029632 | Lrat |
| 97 | PZ81R | Mus musculus | ENSMUSG00000026473 | Glul |
| 98 | PZ85W | Mus musculus | ENSMUSG00000021109 | Hif1a |
| 99 | PZCAY | Mus musculus | ENSMUSG00000030170 | Wnt5b |
| 100 | PZCCZ | Mus musculus | ENSMUSG00000022996 | Wnt10b |
