## Supplementary table for "Dynamic control of metabolic zonation and liver repair by endothelial cell Wnt2 and Wnt9b revealed by single cell spatial transcriptomics using Molecular Cartography": Table S4 .docx

**Table S4 Sequence of qPCR primers** (related to STAR Methods)

| Gene | Forward | Reverse |
| --- | --- | --- |
| *Rn18s* | GTAACCCGTTGAACCCCATT | CCATCCAATCGGTAGTAGCG |
| *Glul* | CTCGCTCTCCTGACCTGTTC | TTCAAGTGGGAACTTGCTGA |
| *Cyp2e1* | AATGGACCTACCTGGAAGGAC | CCTCTGGATCCGGCTCTCATT |
| *Axin2* | TGACTCTCCTTCCAGATCCCA | TGCCCACACTAGGCTGACA |
| *Lect2* | CCCACAACAATCCTCATTTCA | GTTAGCCCATGGTCCTGCTA |
| *Oat* | CCGACCAGTTATGATGGCTTTGG | CTCCACCATGAAGGCAGCAACA |
| *Rgn* | GTATGGGAGGAAGCGTCACAGT | CAATGGTGGCAACATAGCCTCC |
| *Cldn2* | GCAAACAGGCTCCGAAGATACT | GAGATGATGCCCAAGTACAGAG |
| *Gstm1* | ATACTGGGATACTGGAACGTCC | AGTCAGGGTTGTAACAGAGCAT |
| *Lgr5* | CCTACTCGAAGACTTACCCAGT | GCATTGGGGTGAATGATAGCA |
| *Tbx3* | ACTCGGGGTCGGAACTGAA | GGAGGGGGCGATTTTGTTTTT |
| *G6pc* | CAGTGGTCGGAGACTGGTTC | TATAGGCACGGAGCTGTTGC |
| *Pck1* | TGTCTTCACTGAGGTGCCAG | CTGGATGAAGTTTGATGCCC |
| *Arg1* | ACAAGACAGGGCTCCTTTCAG | TGAGTTCCGAAGCAAGCCAA |
| *Cps* | AGGATGTCAAGGTGTTTGGC | GCTTAACTAGCAGGCGGATG |
| *Ass1* | ACACCTCCTGCATCCTCGT | GCTCACATCCTCAATGAACACCT |
| *Atp5a1* | TGGTGAAGAGACTGACGGATGC | TCAAAGCGTGCTTGCCGTTGTC |
| *Igf1* | TCATGTCGTCTTCACACCTCTTCT | CCACACACGAACTGAAGAGCAT |
| *C8b* | ACTGTCAACGGGAGATGGAGCA | GTTGGTGTCCAGGATGTAGTGG |
| *Cyp8b1* | AGTACACATGGACCCCGACATC | GGGTGCCATCCGGGTTGAG |
| *Hpd* | CGCTCCATTGTGGTGACCAACT | TCCGTCTTGAGAGCGATGTGCT |
| *Ccnd1* | TTTCTTTCCAGAGTCATCAAGTGT | TGACTCCAGAAGGGCTTCAA |
